# A rigorous method for integrating multiple heterogeneous databases in genetic studies

**DOI:** 10.1101/2020.09.15.298505

**Authors:** József Bukszár, Edwin JCG van den Oord

**Author notes:** Correspondence should be addressed to Edwin JCG van den Oord.

## Abstract

The large number of existing databases provides a freely available independent source of information with a considerable potential to increase the likelihood of identifying genes for complex diseases. We developed a flexible framework for integrating such heterogeneous databases into novel large scale genetic studies and implemented the methods in a freely-available, user-friendly R package called MIND. For each marker, MIND computes the posterior probability that the marker has effect in the novel data collection based on the information in all available data. MIND 1) relies on a very general model, 2) is based on the mathematical formulas that provide us with the exact value of the posterior probability, and 3) has good estimation properties because of its very efficient parameterization. For an existing data set, only the ranks of the markers are needed, where ties among the ranks are allowed. Through simulations, cross-validation analyses involving 18 GWAS, and an independent replication study of 6,544 SNPs in 6,298 samples we show that MIND 1) is accurate, 2) outperforms marker selection for follow up studies based on *p*-values, and 3) identifies effects that would otherwise require replication of over 20 times as many markers.

**AUTHOR SUMMARY:** The large number of existing databases provides a freely available independent source of information with a considerable potential to increase the likelihood of identifying genes for complex diseases. We developed a flexible framework for integrating such heterogeneous databases into novel large scale genetic studies and implemented the methods in a freely-available, user-friendly R package called MIND. For each marker, MIND computes an estimate of the (posterior) probability that the marker has effect in the novel data collection based on the information in all available data. For an existing data set, only the ranks of the markers are needed to be known, where ties among the ranks are allowed. MIND 1) relies on a realistic model that takes confounding effects into account, 2) is based on the mathematical formulas that provide us with the exact value of the posterior probability, and 3) has good estimation properties because of its very efficient parameterization. Simulation, validation, and a replication study in independent samples show that MIND is accurate and greatly outperforms marker selection without using existing data sets.

## INTRODUCTION

During the past decade, databases related to the genetic basis of complex diseases have grown dramatically. Typical examples are gene expression data repositories, meta-analyses of genomewide linkage scans, published candidate gene association studies, disease-specific biochemical pathways, and genome-wide association studies (GWAS). These databases provide a freely available independent source of information. Integrating this information in novel studies has great potential to increase the likelihood of identifying disease genes. First, the use of existing information may increase statistical power and reduce the risk of false discoveries through improving the prior probability that a marker is associated with the disease. Second, integrating data generated by other technologies may also reduce platform-specific errors and increase confidence in the robustness of the findings when multiple lines of evidence point to the same association. Third, because data integration considers multiple data sources, it may improve the understanding of disease mechanisms by informing the broader context in which disease genes operate^1^.

Data integration may be particularly critical in large scale genetic studies of complex diseases. The reason is that rather than a few markers with large effects, many markers with small effects may be involved. Large sample sizes may therefore be required to find true positives while controlling false discoveries where the cost per sample in these high dimensional investigations is typically high. As a result, economic feasibility may interfere with designing adequately powered studies. Furthermore, with the exception of traits that are routinely measured in control groups of genetic studies (e.g. smoking^2-4^), for many disorders and outcomes studied (e.g. drug response) very large sample size may simply not be available. The use of existing information may then become the only readily available option to detect small effects in a cost-efficient manner.

Because of their volume and heterogeneity, information from existing databases can no longer be integrated intuitively by investigators. This explains efforts towards developing more systematic data integration methods^5-12^. One limitation of these methods is that they lack a solid statistical basis. For example, most methods produce a cumulative measure of the biological relevance of genes after combining information across multiple sources. However, because it is usually very hard to asses the quality of that overall score, it is unclear how to use these scores in a way that information from different databases is used and combined optimally.

In this article we present a rigorous and flexible framework for integrating multiple heterogeneous existing data sets into novel studies aimed at identifying genes affecting complex diseases. We implemented the method in a freely-available R package called MIND (<u>M</u>athematically-based <u>I</u>ntegration of heteroge<u>N</u>eous <u>D</u>ata), that allows researchers to perform all analyses discussed in this paper through a single command line with 8 parameters. MIND can integrate existing data sets generated by any kind of technology (e.g., expression arrays, proteomics, GWAS) or activity (e.g., actual data collection, literature search, construction of disease-specific biochemical networks). Furthermore, external data may provide information at any genetic level ranging from individual variants (e.g., SNPs), genes (e.g., literature search), groups of genes (e.g., pathways), or entire chromosomal segments (e.g., linkage studies or targeted next-generation sequencing).

The end product of MIND is an estimate of the compound local true discovery rate (cℓTDR), which is the posterior probability that a genetic marker has an effect based on the information in the novel data collection and the existing data sets. The adjective “compound” indicates that the cℓTDR capitalizes on (i.e. compounds) all disease relevant information present in the external data sets. The cℓTDR is a posterior probability because it also takes the results from the novel data collection into account. Finally, the term “local” reflects that the cℓTDR provides marker specific information. In scenarios where researchers are interested in groups of markers (e.g. pathways, top results) the cℓTDRs can simply be summed across all markers. The resulting cumulative cℓTDR is then the expected number of markers with effect in that group.

A very important feature of MIND is that, accurately modeling the data integration process and properties of data bases (e.g. number of effects differs across data sets), it relies on a solid mathematical foundation that provide us with the exact posterior probability that a marker has effect in the novel data collection based on the information in all available data sets. This ensures that MIND is more accurate than any heuristic or ad-hoc method. Furthermore, while the exact value of cℓTDR depends on many unknown parameters, most of the unknows can be collapsed into a single parameter for which we developed a precise estimator. As a result, the deviation of the estimate of the cℓTDR from its real value is only due to sample fluctuation, which was also verified by simulation.

A noteworthy property of MIND is that it merely requires that the information in existing data sets can be ranked. This ensures general applicability as ranks can almost always be calculated. For example, one could count the number of times genes co-occur with a specific disease in the literature or use only two ranks indicating whether a gene is implicated or not. Ranks also provide a robust method if there are concerns about the distributional assumptions of the test statistics in the external data sets.

We demonstrate MIND through simulations, cross-validation analyses involving 18 GWAS, and an independent replication study of 6,544 SNPs in 6,298 samples. The markers included in the replication study were selected based on a meta-analysis of the 18 GWAS. The results obtained with the markers selected by MIND are compared with a traditional *p*-value based SNP selection.

## METHODS

The goal of MIND is to identify markers that are associated with a complex disease based on the test statistic values in the novel data collection (NDC) and on their ranks in the existing data sets (EDSs). **Figure 1** displays a schematic overview of the preparatory data transformation as well as the three major steps of the method. First, the existing data sets need to be transformed to the genetic level of the novel data collection if needed (e.g. assign the rank of a gene after an existing literature search to each SNP in that gene in the novel GWAS). The three major steps are 1) Compute for each existing data set the prior probabilities that markers are associated with the disease, 2) Combine the individual sets of prior probabilities into a single set of prior probabilities, and 3) Compute the cℓTDR for each marker.

**Figure 1.**
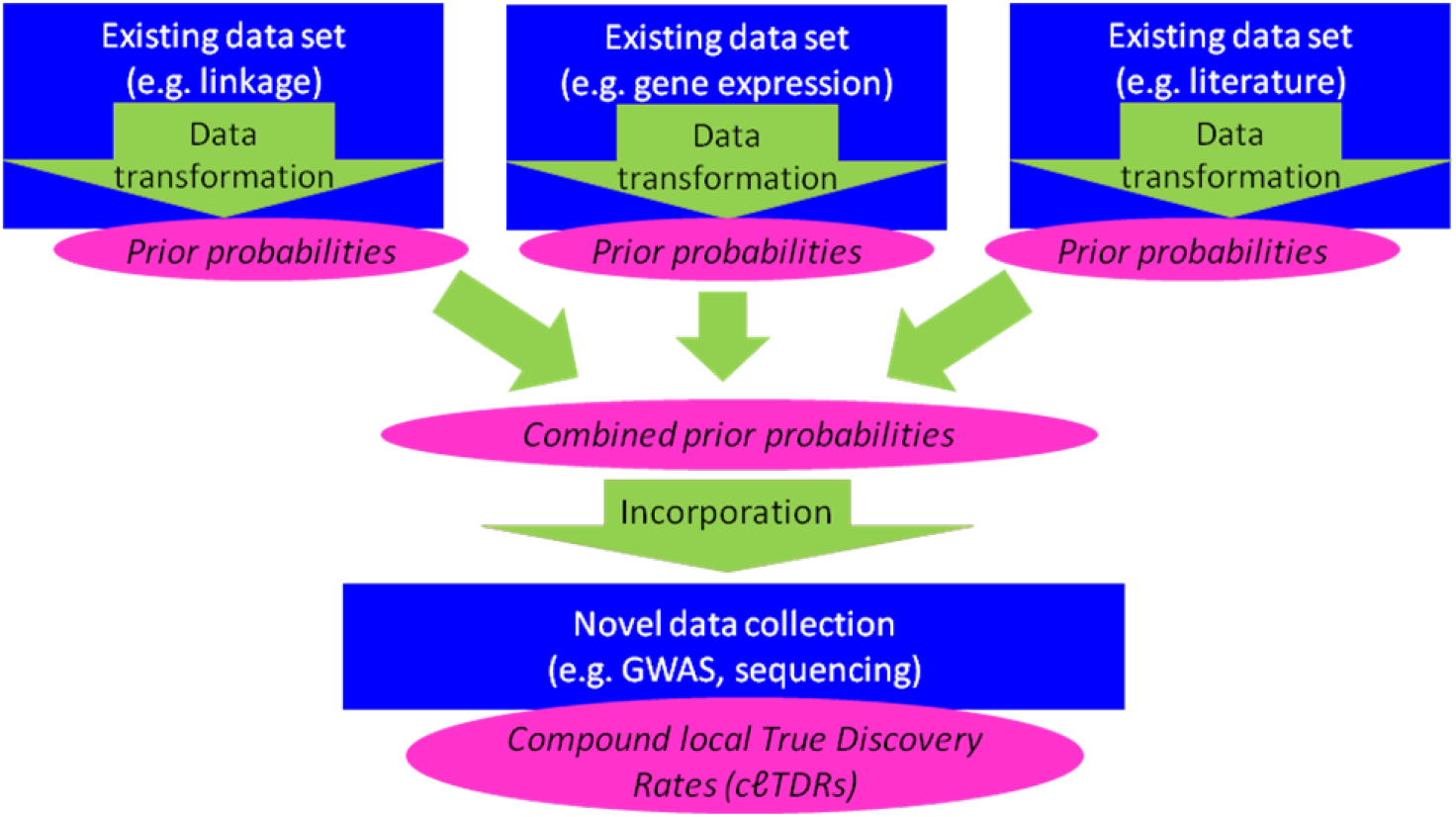
A schematic overview of the MIND data integration framework.

Before discussing each of these three steps, we note for every EDS we only need the rank of their genetic units, whereas for the NDC we need the null and the alternative p.d.f. of the test statistics, *f*_0_ and *f*_1_, as well as the number of alternative genetic units in the NDC, m_1_^**^, or the estimates of them. Although estimating m_1_^**^, *f*_0_ and *f*_1_ is not part of our framework, we developed the estimators of m_1_^**^, *f*_0_ and *f*_1_ for the scenario where the test statistic is approximately normally distributed or its distribution is a mixture of normal distributions in the NDC. Furthermore, MIND allows genetic units to be different across the data sets involved. For instance, if we have gene expression data, GWAS and linkage data as EDS, their genetic units are gene, SNP, and chromosomal segment, respectively. To handle these different units we first transform the EDSs into data sets based on the genetic unit of the NDC, which we call the test unit. For instance, a gene-based EDS can be transformed into SNP-based EDS by assigning to each SNP the smallest EDS rank (or p-value) of the genes that contain the SNP.

In the rest of this section we describe the three major steps. For each step we also present the mathematical formula based on which the step is carried out. The mathematical proofs for the formulas are provided in the Supplemental Material, Appendix.

<u>Step 1: Obtaining prior probabilities of test units for each EDS</u>: First we need two concepts to quantify information. We define the *information parameter* of an EDS to the NDC as

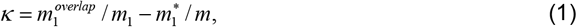

where *m* denotes the number of test units that are both in the EDS and in the NDC, *m_1_* is the number of test units alternative in the EDS, *m_1_^*^* is the number of test units in the EDS that are alternative in the NDC, and *m_1_^overlap^* is the number of test units that are alternative in the EDS and in the NDC. We will call an EDS *informative to* the NDC if its information parameter is positive. Note that κ is positive if, and only if, the number of test units alternative both in the NDC and the EDS is larger than it would by chance, i.e. when the alternative label would be randomly assigned to the test units of the EDS.

For test unit *i* in an EDS, we define *the contribution of test unit i from the EDS to the NDC* as

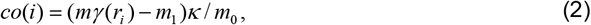

where *r_i_* is the rank of test unit *i* in the EDS, *γ*(*r*) denotes the probability that a test unit ranked *r* in the EDS is alternative in the EDS, and *m_0_=m-m_1_*.

Based on the information in an EDS, for the prior probability that test unit *i* is alternative in the NDC, *γ^*^*(*i*), we have that

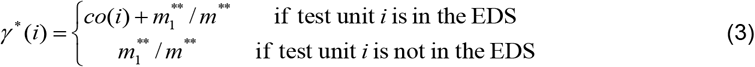

where *co*(*i*) is the contribution of test unit *i* from the EDS to the NDC, *m^**^* and *m_1_^**^* is the number of test units and the number of alternative test units in the NDC, respectively (for the proof see Theorem 3 and Corollary 4 in **Appendix**). We remark that the information parameter being 0 implies 0 contributions for all test units, which results in *γ^*^*(*i*)= *m_1_^**^/m^**^* for every test unit. Note that this is exactly what we would have in case of no prior information. Another property of the *γ^*^*(*i*) formula is that, as all the contributions of an EDS sum up to zero on the test units of the EDS (see Lemma 8 in **Appendix**), *γ^*^*(*i*) sum up to *m_1_^**^* on the test units for every EDS. In other words, using an EDS merely redistributes the total amount of prior probabilities among the test units.

The contribution of a test unit depends on 3 factors: 1) the rank of the test unit in the EDS, 2) the information parameter of the EDS and 3) the effect sizes in the EDS (see (2)). These latter two, intuitively speaking, stretch out the contributions, and hence amplify the redistribution of the prior probabilities. Indeed, larger information parameter or average effect size of the EDS make the contributions differ from each other more within the EDS. It is an advantage of our method, however, that we do not need to know the information parameter and the average effect size separately to obtain the prior probabilities, because only their combined effects matter, which we can estimate from the data.

To obtain (3) we utilized that in practice, we can approximate *m_1_^*^/m* by *m_1_^**^/m^**^*, where the rationale is that the group of test units in the NDC we have EDS information for should contain proportionally as many alternatives as the entire NDC does (see Corollary 4 in **Appendix**). On the other hand, if we think that the above assumption is violated, then we may be able to estimate *m_1_\** in the same way as *m_1_\*\** is estimated, and use Theorem 3 to obtain the prior probability estimates.

<u>Step 2: Combining the sets of prior probabilities into a single set of prior probabilities</u>: Once we have the prior probabilities for every EDS, we calculate the combined prior odd 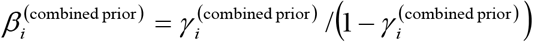 that test unit *i* is alternative in the NDC by

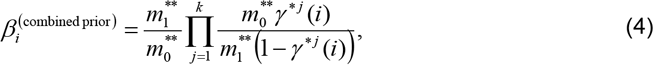

where *γ^*j^*(*i*) is the *j*th EDS-based prior probability that test unit *i* is alternative in the NDC, and *m_0_^**^ = m^**^ – m_1_^**^* (see eq. 13 in **Appendix**). Note that if we have no prior information for a test unit in any EDS, then from the formula in (4) we obtain that the combined odd of this test unit is *m_1_^**^/m_0_^**^*, which is exactly what we supposed to have in the case of no prior information. Moreover, according to the formula in (4), the combined odd of a test unit is proportional to the average of the prior odds of the test unit across the EDSs, where by the average we mean the geometrical mean. If a test unit performs better than a test unit with no information in an EDS (odd = *m_1_^**^/m_0_^**^*), then its odd in that EDS will have a positive (increasing) impact on its combined odd, and vice versa, i.e. if a test unit performs worse than a test unit with no information in an EDS, then its odd in that EDS will have a negative (decreasing) impact on its combined odd.

<u>Step 3: Computing cℓTDR for each test unit</u>: The cℓTDR of test unit *i* can be written as

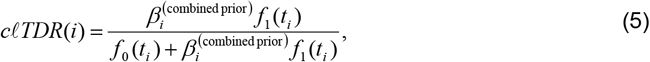

where f_0_ and f_1_ is the null and alternative p.d.f. in the NDC, respectively, and where *t_i_* is the observed test statistic value of test unit *i* in the NDC (see Claim 24 in **Appendix**). In summary, combining equations in (3), (4) and (5) we obtain the cℓTDR of a test unit as a function of *m_1_^**^*, f_0_, f_1_, and the contributions of test units from each EDS. Instead of the terms cℓTDR depends on, we will use their estimates to obtain estimate of the cℓTDR. In the next section we present a method that estimates the contributions.

### Estimating the contributions

As mentioned above, in order to estimate cℓTDR by our formulas we need to estimate the contributions of test units from an EDS to the NDC, defined in (2). Because we use the same procedure to estimate contributions for each EDS, throughout this subsection we assume that we have a single EDS, which we will refer to as the EDS. As we focus on the test units in the NDC, it is irrelevant whether the EDS contains test units not in the NDC or not, so for the sake of simplicity, we assume that the test units the EDS contains are also in the NDC. For estimating the contributions we will use the statistic

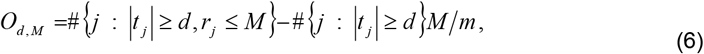

where #*A* denotes the number of elements in set *A, t_j_* is the NDC test statistic of test unit *j*, and *r_j_* is the rank of test unit *j* in the EDS. In **Appendix** (Theorem 25) we proved that for any positive integer *M ≤ m* and real number *d* ≥ 0 we have that

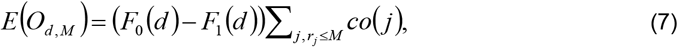

where *F*_0_ and *F*_1_ is the null and alternative c.d.f. in the NDC, respectively. Based on eq (7), first we calculate a rough estimate of the cumulative contribution, defined as 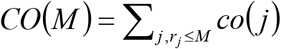 by

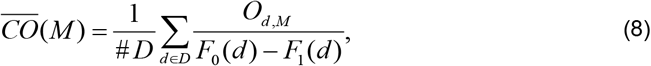

where *D* is a set of the positive real numbers, and #*D* denotes the number of elements in *D*. As the contribution of the test unit whose rank is *r* in the EDS can be obtained as

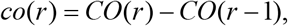

we can calculate the contribution estimates from estimates of the cumulative contribution. To ensure that test units with smaller (better) ranks have larger contribution estimates, we need to use a cumulative estimate that is a concave function of M. For this we construct a concave function of M that fits 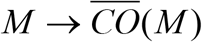 well (see Section 1.3 in **Appendix** for details).

## RESULTS

### Accuracy

To study the accuracy of MIND, we simulated 500 studies with 3 existing data sets and a novel data collection consisting of one million markers of which 5,500 had a small effect (see **Supplemental Material** for details). In **Figure 2**, we show the estimates of the cumulative cℓTDR, defined as the sum of the *k* largest cℓTDRs as a function of *k*. The cumulative cℓTDR at *k* equals the expected number of markers with effect among the *k* markers with the largest cℓTDRs. For the sake of comparison, in the figure we also show the corresponding curves where no existing data sets were used to compute the cℓTDR estimates. Each curve in Figure 2 is the average of the corresponding curves in the 500 simulation studies. The fact that the lines overlap perfectly implies that on average the estimated cumulative cℓTDRs are very precise indicators of the number of markers with effect in the novel data. Moreover, we found that the cumulative cℓTDR differs from the number of markers with effect among the markers with largest cℓTDRs by less than 9.6% of the number of markers with effect among the markers with largest cℓTDRs in 99% of the simulations studies, and the percentage difference gets smaller as the number of selected markers increases (see **Supplemental Material** for details). This shows that the estimated cumulative cℓTDR is an accurate predictor of the number of markers with effect among the markers with largest cℓTDRs. Comparing in **Figure 2** results for the cℓTDR when the existing data sets were used versus when no existing data sets were used shows how data integration increases the proportion of markers with effect among markers selected by their cℓTDR.

**Figure 2.**
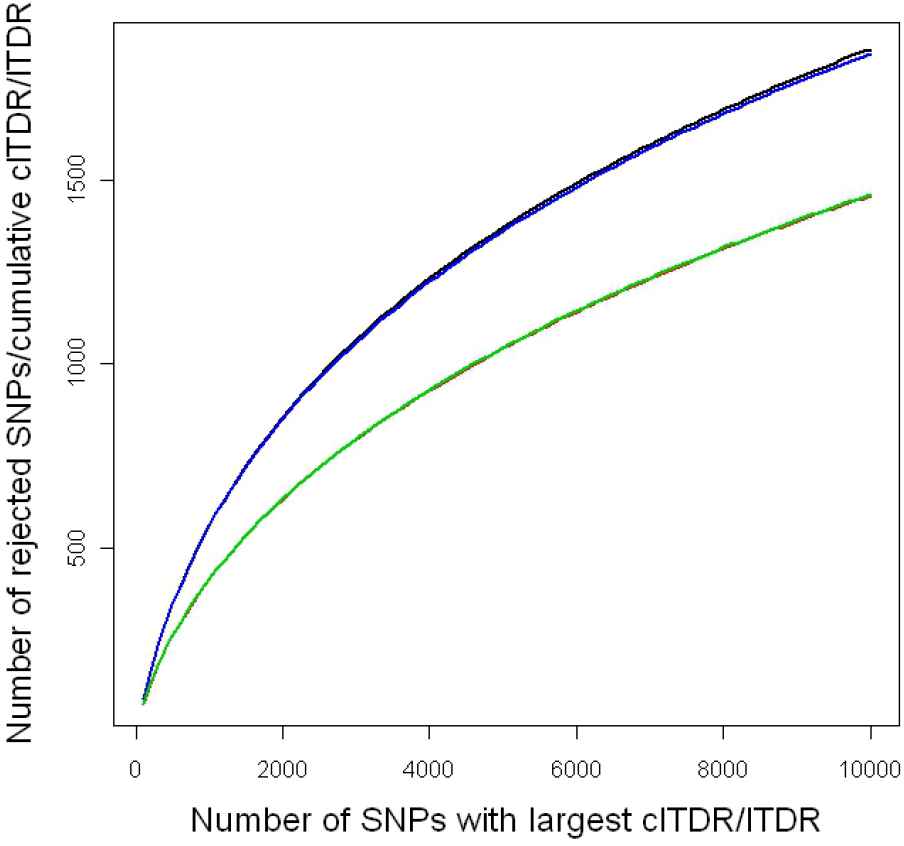
The estimated cumulative cℓTDR (black) as well as the number of markers with effect among the markers with the largest cℓTDRs (blue) curves are plotted. We also plotted the corresponding curves, where no existing data sets were used to compute the cℓTDR estimates (red and green). Each curve is the average of the corresponding curves in the 500 simulation studies.

### Illustration with empirical data

We illustrate our framework using a meta-analysis of 18 schizophrenia GWAS studies comprising a total of 21,953 cases and controls. Even after including study-specific principal components to control for stratification, the meta-analyses suggested the presence of many SNPs with very small effects (consistent with a previous publication^13^). To distinguish and select these very small effects from the markers with no effects, we used MIND. Nine external data sets with potential relevance for schizophrenia were tested for information content 1) schizophrenia candidate genes^14^, 2) the top bins from a meta-analysis of linkage scans^15^, 3) results from an expression array meta-analysis using post-mortem brain tissue from schizophrenia cases^16^, 4) a global proteomic analysis in post-mortem prefrontal brain tissues^17^, 5) CNVs associated with schizophenia^18^, 6) disease genes in the OMIM database^19^, 7) gene length, 8) gene expression quantitative trait loci (eQTLs)^20-25^, and 9) human orthologs of murine genes showing association with behavioral phenotypes relevant to neuropsychiatric outcomes^26^. Six (above data sets 1, 2, 3, 6, 8, and 9) out of these 9 external data sets appeared informative for the GWAS meta-analyses and were included in subsequent analyses. We note that our finding that eQTL data are informative for GWAS is consistent with other reports in the literature^27^.

In **Figure 3** we use the (meta-analyses of) expression data to graphically illustrate informativeness. To each GWAS SNP that was ±50kb of a gene in the eQTL dataset, we assigned the rank of the *p*-value of that gene in the expression data. We picked the smallest *p*-value if there were multiple *p*-values per gene. A total of 441,392 GWAS SNPs could be assigned a rank. The x-axis shows the top *j* SNPs according to their rank in the expression data set. The purple line gives the relation as observed in the data, and the many thin grey lines show results from 1,000 generated existing data sets obtained by randomly permuting the ranks of genes in the expression data set. The figure shows that up to about the first 40,000 SNPs, SNPs that are in genes that rank higher in the expression data also have better *p*-values in the GWAS and that this pattern is unlikely to occur by chance.

**Figure 3.**
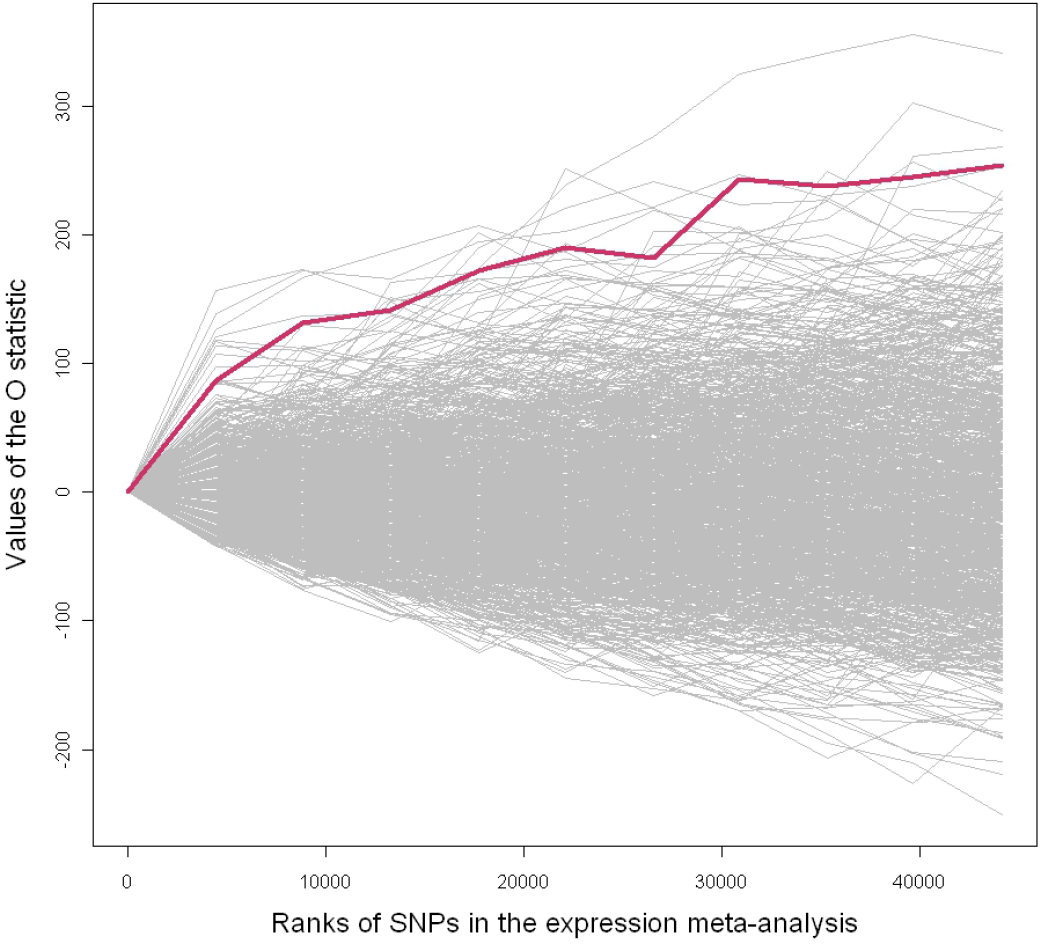
Enrichment in GWAS *p*-values (y-axis) in the set of SNPs that are in genes with higher ranks in the expression data. The purple line represents the expression data set, and each of the thin grey lines shows results from 1,000 data sets generated by random permutation.

As an initial “internal” validation, we compared the 5,000 SNPs with the best *p*-values versus the 5,000 SNPs with the best cℓTDRs in terms of the heterogeneity of effects and gene ontology (see **Supplemental Material).** The *I*^2^ index^28^ was used to study heterogeneity that could, for example, be increased due to technical errors affecting results from only one or a few GWAS. Integrating data generated by other technologies may reduce the effect of such technical errors. We therefore hypothesized that compared to *p*-value selected SNPs, the cℓTDR selects SNPs that show more consistent effects across the 18 GWAS in our meta-analysis. Results in **Supplemental Figure 2** confirm that this is indeed the case. The gene ontology analysis is motivated by the observation that most biological functions seem to be carried out by coregulated “modules” (e.g. pathways, complexes)^29^. Some of these modules could possibly be pathogenic, implying that disease genes may share gene ontology terms. For this specific analysis we only integrated the empirical external data sets (e.g. linkage analyses, expression array) to avoid that differences were introduced by using databases that we (partially) generated using biological knowledge (e.g. candidate gene studies). Results (see **Supplemental Figure 3**) support for the notion that data integration more successfully identifies gene ontology terms thereby improving our understanding of disease mechanisms.

### Validation

We validated the ability of our data integration method to identify markers with effects via 1) simulation studies, 2) cross-validation, and 3) and actual replication study of 6,544 SNPs in a sample independent of the 18 GWAS studies that included 6,298 subjects from 1,811 nuclear families. In the simulation studies we used the same parameters as used in the accuracy section above. For cross-validation, presenting a more realistic test case (e.g. actual effects sizes, artifacts, LD among markers), we selected subsets from all 18 GWAS in such a way that the sample size available for selecting SNPs was 85-90% of the total sample size. The remaining studies were used for replication/cross-validation. For each of the 575 unique cross-validation combinations, we selected the 5,000 SNPs with the smallest *p*-values and the 5,000 SNPs with the best cℓTDR after integrating our six informative existing data sets. The replication study involving genotyping of 6,544 SNPs in independent samples was conducted using a custom Illumina iSelect chip. About half of the SNPs were selected based on having the smallest *p*-values and the other half based on having the best cℓTDRs.

Results are shown in **Figure 4a, b**, and **c**. All three panels converge to the same conclusions. First, considering the cℓTDR (green and blue dots) always gives better results compared to SNP selection based on *p*-values alone. Second, although the success of MIND decreases as *p*-values increase, in many instances it still successfully identifies SNPs with effects that have large *p*-values with ranks >100,000 in the GWAS meta-analysis. Even if costs to follow up that many SNPs would not be an issue, it may still not produce equally good results because, compared to the much smaller set of cℓTDR selected SNPs, many more tests would need to be performed in the replication study. Third, these cℓTDR selected SNPs with *p*-value ranks >100,000 in the meta-analysis replicate as well as or better than SNPs with small *p*-values but poor cℓTDRs.

**Figure 4.**
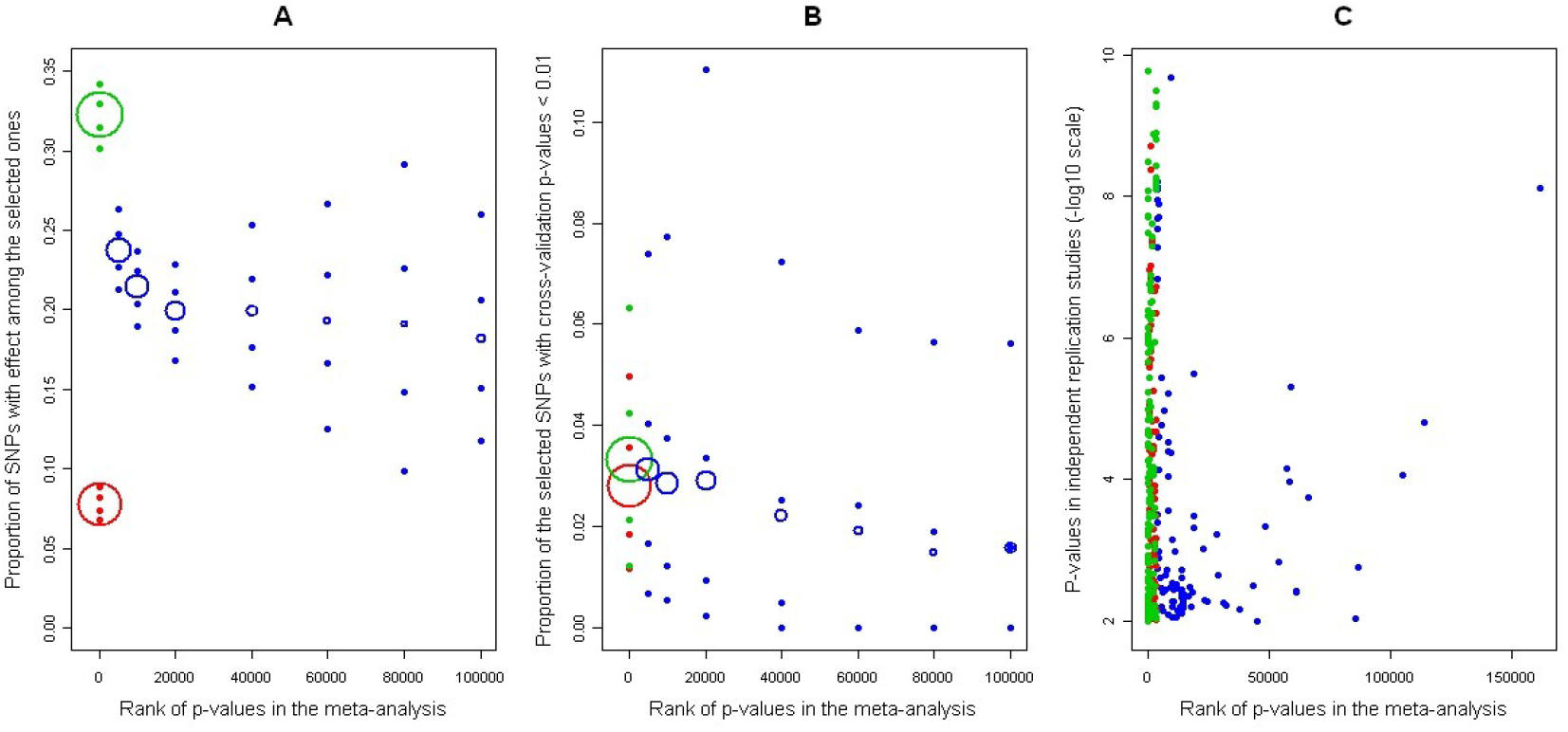
Performance of *p*-value versus cℓTDR based selection by simulation (A), crossvalidation (B), and replication (C). All panels: *p*-value based results are red, cℓTDR based results are blue, overlap between both methods is green, and x-axis is rank of p-values in the metaanalysis. <u>Panels A and B</u>: For each x-axis interval, the .05, .25, .75 and .95 quantiles of the proportion of SNPs with effect among those selected (A) or proportion of SNPs with crossvalidation *p*-value less than 0.01 among those selected (B) are reported. Center of the circles are located at the mean of the proportions, while the area of the circle is proportional to the number of SNPs selected. <u>Panel</u> C: all *p*-values less than 0.01 in the replication study are shown.

## DISCUSSION

We developed a mathematically rigorous and flexible framework for integrating heterogeneous databases into large-scale genetic studies, and implemented our method in a freely available user-friendly R package called MIND. Through simulations, cross-validation analyses involving 18 GWAS, and independent replication of 6,544 SNPs in 6,298 samples we show that MIND 1) is accurate, 2) outperforms marker selection for follow up studies based on *p*-values, and 3) is able to identify effects that would otherwise require replicating over 20 times more markers.

Although a main application of MIND involves integrating existing data in a novel data collection, it is applicable in other scenarios as well. For example, MIND can rank genes according to their relevance to a disease using only existing databases. However, it is typically difficult to assess the quality of such prioritization scores; our framework provides an estimate of the probability that a gene is associated with the disease of interest. This clear interpretation allows for more informed decisions about which genes to select for further study. A second example involves questions related to similarities among multiple high dimensional data sets. For example, if we have datasets for different diseases in the same population, MIND can be used to study co-morbidity where a high concordance would indicate a substantial overlap in disease etiology. Alternatively, if we have datasets for the same disease in different populations, MIND would shed light on the overlap in the genetic disease architecture of the different populations.

We should stress that MIND can handle novel data collections of any kind. Nextgeneration DNA sequencing (NGS) has the potential to accelerate genetic research. However, because costs for NGS are still high and power to detect the (cumulative) effects of all rare variants low^30^, data integration could play an important role. Indeed, several methods have already been proposed that test for association using weights based on predictions of functional effects of (rare) variants (e.g. ^31-32^). However, as these weights do not take the strength of disease relevant information into account, our method could be used to further optimize these tests. A second example is that NGS enables a comprehensive analysis of not just genomes but also transcriptomes and methylomes. MIND offers the possibility to integrate all these different sources of information to improve statistical power, increase confidence in the robustness of the findings when multiple lines of evidence converge to the same genetic factors, and inform the broader context in which the disease genes operate.

## SOFTWARE

MIND has been made freely available as an R package at http://www.people.vcu.edu/~jbukszar/.

## ACKNOWLEDGMENTS

J.B. and EvdO are supported by grants R01HG004240; R01MH078069; and HG004240-02S1.

## Supplemental material

### 1. Simulation study

In order to study how accurately the cumulative cℓTDR predicts the number of selected SNPs that have effect in the novel data collection, we used 500 simulations. In each simulation, we generated 3 existing data sets and a novel data collection. For the novel data collection we simulated 1,000,000 test statistic values, 5,500 of which had effect. To generate an existing data set we randomly chose 500,000 of the 1,000,000 NDC SNPs to be matched and simulated test statistic values for them, 50,000 of which had effect. The “alternative in the EDS” label was randomly assigned to SNPs in such a way that the number of SNPs alternative both in the NDC and EDS was 2,200. We calculated the ranks of the test statistics in the EDSs, which were used for the data integration. The statistic values were drawn from the normal distribution with variance 1. The mean of the normal distribution was 1.6 and 2.0 for the SNPs with effect in the existing data sets and the novel data collection, respectively, and the mean was 0 for the null SNPs for every data set. As estimating the null and alternative distribution of the statistics as well as the number of SNPs with effect in the novel data collection is not part of our method, we used the ‘real’ functions and number of SNPs with effect for our procedures.

The average of estimated cumulative cℓTDR (=after data integration) and ℓ TDR (=before data integration) curves as well as the number of SNPs with effect in the NDC in the top SNPs for cℓTDR and ℓ TDR – based selection are plotted in the left panel of Figure S1 (identical to Figure 1 in the article). Each curve is the average of the corresponding curves in the 500 simulation studies. The figure suggests that the cumulative cℓTDR and the cumulative ℓTDR curves are unbiased predictors of the number of SNPs with effect in the NDC that were selected by the corresponding method. To further study the accuracy of these predictors, we calculated the percentage difference between the cumulative cℓTDR/ℓTDR and the number of SNPs with effect selected by cℓTDR/ℓTDR, that is the absolute value of the difference between the two expressed with the percentile of the number of SNPs with effect selected by cℓTDR/ℓTDR. The quantiles of the percentage differences in the 500 simulation studies are plotted in the right panel in Figure S1. For instance, the continuous black curve in the figure shows that in 99% of the simulation studies the percentage difference between the cumulative cℓTDR and the number of selected SNPs with effect in the NDC was always less than 9.6%, and less than 7.5% if more than 5,000 SNPs were selected. The 50%, 75%, 90% and 99% quantile curves of the percentage difference between the cumulative cℓTDR/ℓTDR and the actual number of selected test units show that percentage difference 1) gets smaller as the number of selected markers gets larger and 2) is smaller for cℓTDR selection than for ℓTDR selection when the number of selected markers is small and comparable otherwise. We conclude that the estimated cumulative cℓTDR is a good predictor of the number of markers with effect among the selected ones and may be limited only by the imperfection of the estimate of the null and alternative distribution of the statistics and the number of markers with effect in the novel data collection, which estimate is, however, not part of our method.

**Figure s1.**
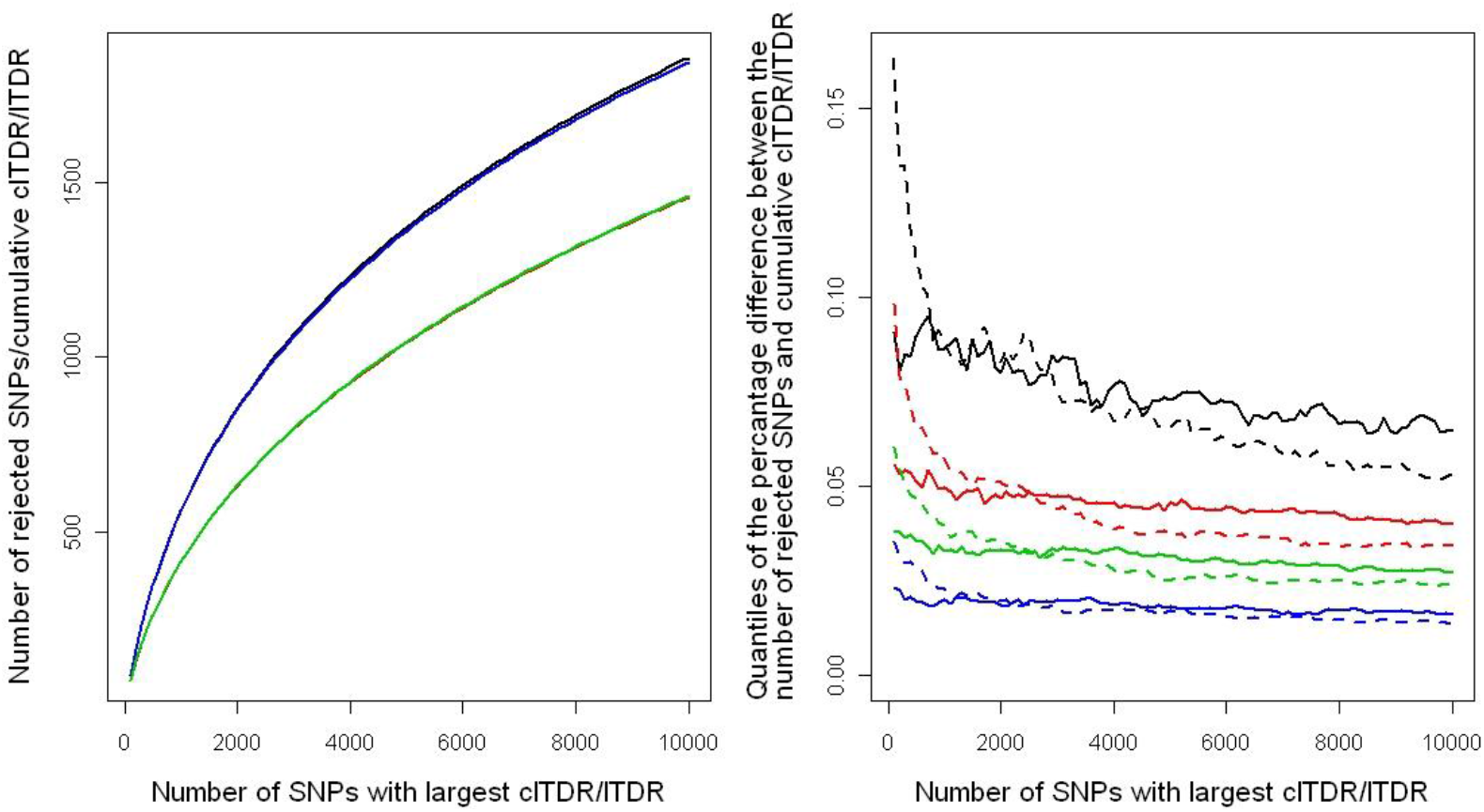
Left panel: The estimated cumulative ℓTDR/cℓTDR (red and black) as well as the number of SNPs with effect among the ℓTDR/ cℓTDR selected ones (blue and green) curves are plotted. Each curve is the average of the corresponding curves in the in the 500 simulation studies. <u>Right panel:</u> Multiple quantiles of the absolute value of percentage difference between the estimated cumulative ℓTDR/ cℓTDR and the number of SNPs with effect among the cℓTDR selected ones are plotted. Black, red, green and blues curves represent the 99%, 90%, 75% and 50% quantiles, respectively, while continuous and dashed curves represent cℓTDR and ℓTDR selection, respectively.

16.4%, 9.9%, 6.1%, respectively, for the 99%, 90% and 75% confidence intervals.

## 2. Data sets and QC

### GWAS meta-analysis

In our empirical example we used a meta-analysis we performed involving 18 schizophrenia GWAS studies. After stringent QC 1,085,772 (imputed) SNPs were available for 21,953 subjects (11,185 cases and 10,768 controls). To account for possible population stratification effects within each of the GWAS studies, we included the first 3 principal components obtained with EigenSoft^14^ plus any additional principal components if they significantly (*p* < 0.05) predicted case-control status.

### External data sets

Our external data sets included 1) schizophrenia candidate genes from the SZgene data base^15^ that summarizes the results of 1,617 studies reporting on 952 candidate genes, 2) the top bins from a meta-analysis of 32 independent genome-wide linkage scans that included 3,255 pedigrees with 7,413 genotyped cases affected (see Table 2)^16^, 3) results from an expression array meta-analysis of 12 controlled studies across 6 different microarray platforms using brain tissue from schizophrenia, bipolar, and controls (about 35 subjects in each group)^17^, 4) a global proteomic analysis in postmortem prefrontal brain tissues of 9 schizophrenic patients and 7 controls^18^, and 5) replicated and significant CNVs (see Table 2^19^) from 10 studies. Other data sets involve features of disease genes in general such as 1) genes present in the OMIM database, 2) gene length, disease genes are suggested to be longer^20^, 3) SNPs that are strongly associated with variation in transcript abundance in the following tissues: liver, cortex and large B-Cell lymphomas using the eQTL browser at U. Chicago^21–26^, and 4) human orthologs of murine genes showing association with behavioral phenotypes relevant to neuropsychiatric outcomes^27^.

## 3. Details results and items in the text

### Test results for informativeness

Test results for informativeness are shown in **Table S1** and indicate that six out of the 11 existing data sets appeared informative for the GWAS meta-analyses. The non-informative existing data sets are the ones with small samples sizes or not directly related to schizophrenia.

**Table S1:** Information tests for existing data sets

| Data type | Source | Unit | Informative? |
| --- | --- | --- | --- |
| Meta-analysis expression studies |  |  |  |
| Schizophrenia | Stanley | Gene | Yes |
| Bipolar | Stanley | Gene | No |
| Candidate genes |  |  |  |
| Schizophrenia | SZgene database | Gene | Yes |
| Bipolar | SLEP + literature | Gene | No |
| Disease genes | OMIM | Gene | Yes |
| Meta-analysis schizophrenia linkage studies | Nga et al. (2009). | Region | Yes |
| Human (neurological) genes with mouse orthologs | SLEP | Gene | Yes |
| Candidate schizophrenia CNV regions | Sebat et al. (2009) | Region | No |
| Gene length | ENSEMBL database | Gene | No |
| Schizophrenia proteomics | de-Souza et al (2009) | Gene | No |
| Expression QTLs | Browser at U. Chicago | SNP | Yes |

### Figure 2

Figure 2 allows a visual inspection of informativeness. The x-axis shows the top *j* SNPs according to their rank in the expression data set. The y-axis shows a measure of enrichment, indicating whether GWAS *p*-values of SNPs are better for the genes that rank higher in the expression data. More precisely, if *p*_i_ indicates the *p*-value of SNP *i* in the GWAS, *ri* indicates the rank of SNP *i* in the expression data, and *j* is an arbitrary a cut-off for the rank of SNPs in the expression data, then enrichment (O statistic) is defined as

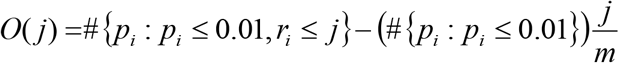

where *m* is the number of SNPs in the expression data. Clearly, *O*(*j*) being positive suggests enrichment of GWAS p-values smaller than 0.01 among the SNPs ranked j or better in the expression data. The purple line in Figure 2 gives the observed relation and the many thin lines show results from 1,000 generated existing data sets obtained by randomly permuting the ranks of genes in the expression data set. The results show that up to the first ~40,000 SNPs, SNPs that are in genes that rank higher in the expression data also have better *p*-values in the GWAS and that this pattern is unlikely to occur by chance.

### Heterogeneity

To study heterogeneity we used the index^4^ *I*^2^ = 100%×(*Q* – df)/*Q*, where (Cochran’s) *Q* is computed by summing the squared deviations of each study’s estimate from the overall meta-analytic estimate, weighting each study’s contribution in the same manner as in the metaanalysis. *I*^2^ describes the percentage of total variation across studies due to heterogeneity rather than chance. Larger values show increasing heterogeneity and the (lower bound) value of 0% indicates no heterogeneity.

**Figure s2.**
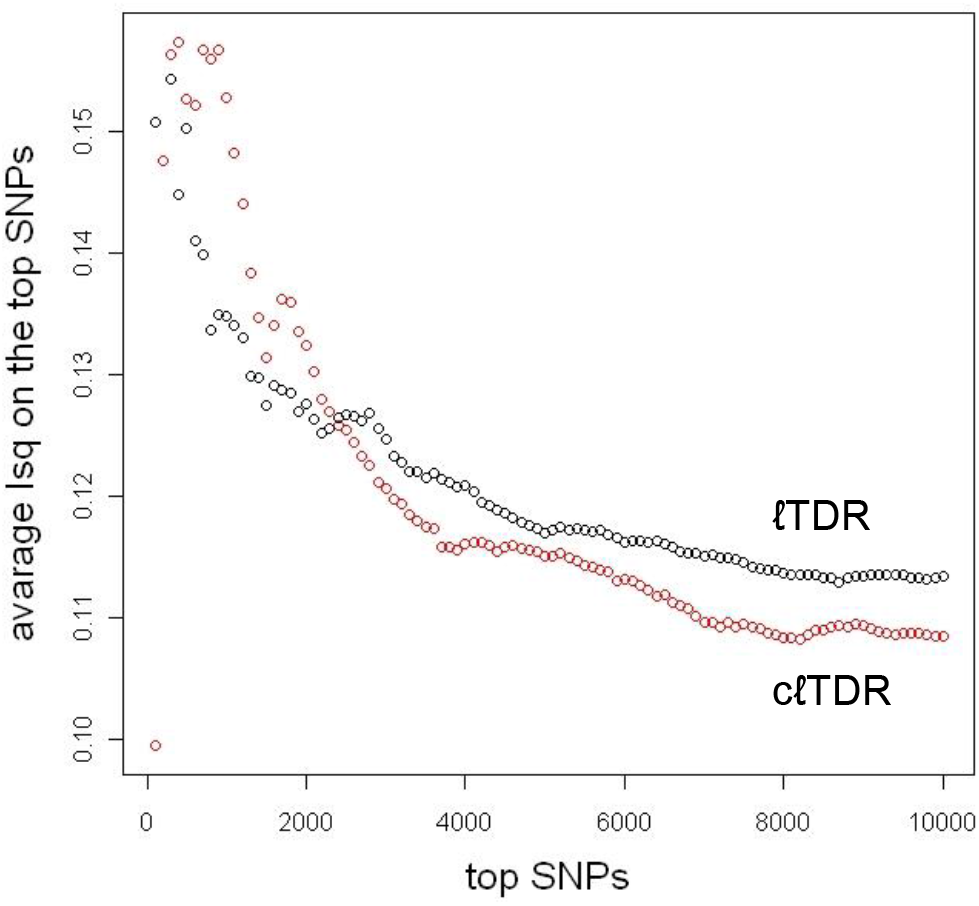
Heterogeneity of SNP effects before and after data integration. The y-axis shows the average *I*^2^ index and the x-axis the top SNPs ranked in ascending order using either the ℓTDR (red) or c.TDR (black).

Results in **Figure S2** show the average *I*^2^ of the top SNPs, with the number of SNPs indicated on the x-axis, selected before and after data integration. As the number of SNPs used to calculate the average *I*^2^ is small at the left hand side of the figure, the results fluctuate initially. However, as the number of SNPs increases, the average *I*^2^ values becomes better for the cℓTDR compared to the ℓTDR, implying that data integration results in the selection of SNPs with more consistent effects across the 18 GWAS studies.

### GO analyses

We performed gene ontology (GO) analyses to establish whether genes selected through data integration show differences in terms of GO themes. For these analyses we only used empirical data sets (linkage analyses, expression array) to avoid that enrichment was introduced by using external data sets that are based on biological knowledge (e.g. candidate gene studies). We first selected SNPs based on having a good (*n* = 1,435) with a larger posterior probability of belonging to the group of SNPs with small effects than the other two groups. Then we identified the genes in which these SNPs were located and subjected those SNPs to a GO analysis to search for biological themes. A similar number of genes were selected using the top ℓTDR results. As an additional control groups, we also selected a similar number of genes from the bottom (i.e. highest values) of the cℓTDR and ℓTDR distributions.

For the GO analyses we used GOEAST^2^, a web based software toolkit. The ontology covers three domains: *cellular component*, the parts of a cell or its extracellular environment; *molecular function*, the elemental activities of a gene product at the molecular level, such as binding or catalysis; and *biological process*, operations or sets of molecular events with a defined beginning and end, pertinent to the functioning of integrated living units: cells, tissues, organs, and organisms. GOEAST uses an exact (hypergeometric) test to evaluate the null hypothesis that genes are picked at random from the total gene population.

**Figure S3** shows the results for the top (blue line) and bottom (green line) genes selected before (**Fig. a**) and after (**Fig. b**) data integration. The y-axis shows the *p*-values and the X-axis shows the number of genes with *p*-values smaller than 0.1 and then sorted in ascending order. Compared to the control group of genes selected from the bottom of the cℓTDR and ℓTDR distributions, there are many more genes with *p*-values smaller than 0.1 and the tests also indication much smaller *p*-values. Furthermore, this pattern is much more pronounced after the data integration. Thus, our top results differ from the bottom results in terms of gene ontology and data integration tends to increase the distinction between top and bottom genes.

**Figure S3.**
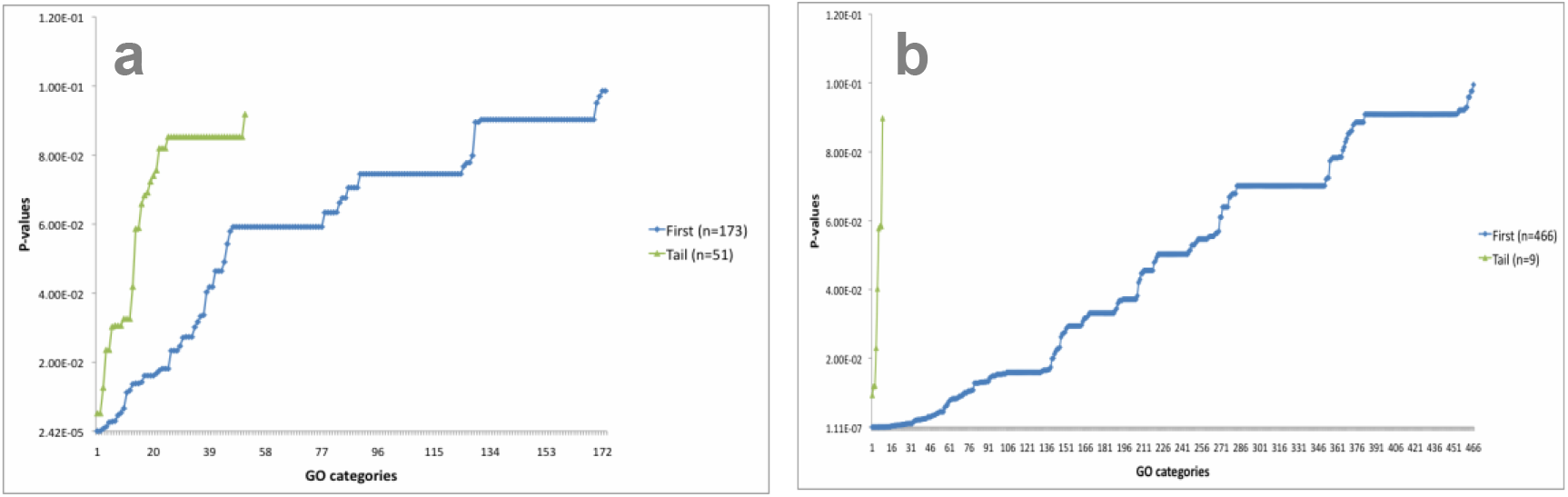
GO analysis on the top (blue line) and bottom (green line) genes selected before (Fig a) and after (Fig b) data integration. The y-axis shows the *p*-values testing the null hypothesis that the selected genes were picked at random from the total gene population and the X-axis shows the number of genes with *p*-values smaller than 0.1 and then sorted in ascending order.

## Appendix

### 1.1 Mathematical formulas for the exact computation of cℓTDR

#### Step 1: Obtaining prior probabilities of test units for each EDS

The goal of this section is to derive the equation of prior probability (Theorem 3). Throughout this section we assume that we have a single existing data set. For brevity we will use the notation *γ_r_* instead of *γ*(*r*) for the probability that a test unit ranked *r* in the existing data set is alternative in the existing data set.

##### Theorem 1

*Suppose we have only one existing data set. Denote the number of test units of NDC that are also in the existing data set as m. Out of these m test units, denote the number of those alternative in the NDC, the existing data set and in both data sets as* 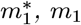, *and* 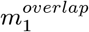, *respectively, and let m*_0_ = *m − m*_1_. *Then we have that*

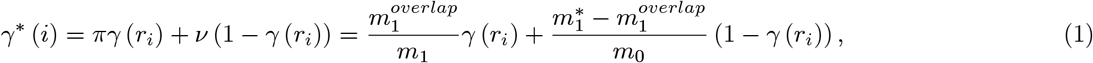

*where r_i_ is the rank of test unit i in the existing data set and*

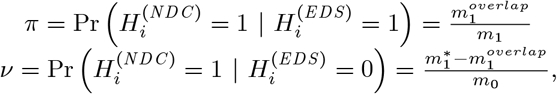

*where* 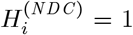 *or* 0 *if test unit i is alternative or null in the NDC, respectively, and* 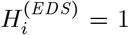 *or* 0 *if test unit i is alternative or null in the existing data set, respectively.*

**Proof.** By definition

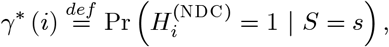

where *S = s* represents our information from the existing data set. Applying the well-known identity Pr (*B | C*) = ∑_*i*_ Pr (*B | A_i_, C*) Pr (*A_i_ | C*), where {*A_i_*} is a partition of the probability space, we obtain that

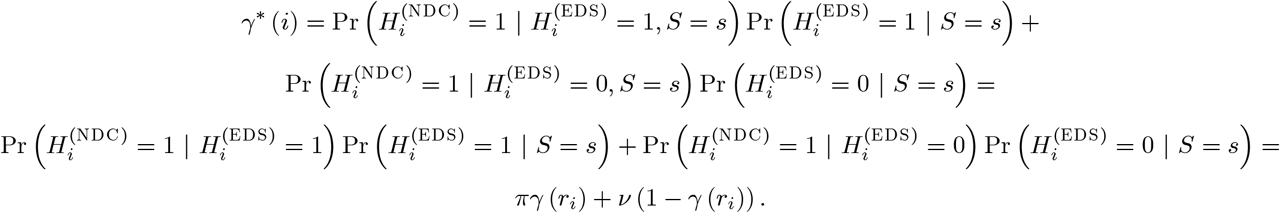

##### Lemma 2

*For test unit i in the existing data set we have that*

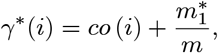

*where*

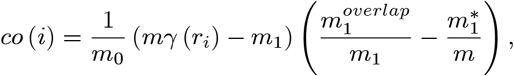

*and r_i_ is the rank of test unit i in the existing data set, m is the number of test units of NDC that are also in the existing data out of which* 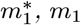, *and* 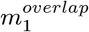, *is the number of test units alternative in the NDC, the existing data set and in both data sets, respectively*.

**Proof.** For brevity we will denote *r_i_* as *j*, hence *γ*(*r_i_*) ≡ *γ*(*j*) ≡ *γ_j_*. From (1) we have that

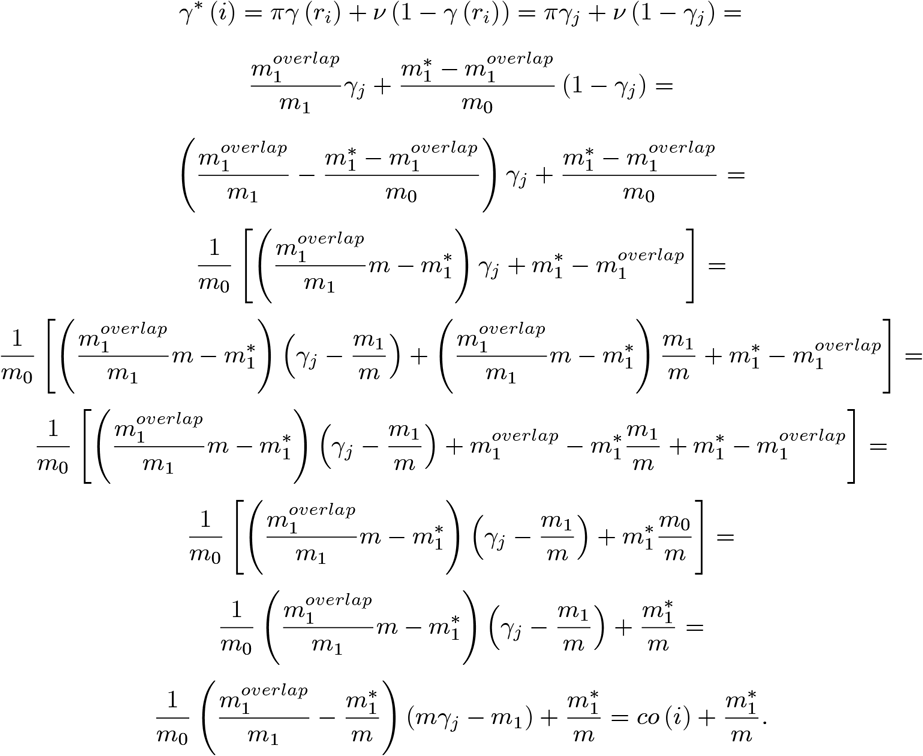

##### Theorem 3

*For test unit i in the NDC we have that*

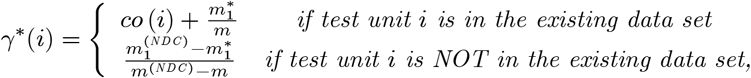

*where m is the number of test units of NDC that are also in the existing data set, out of which* 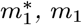, *and* 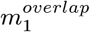 *is the number of test units alternative in the NDC, the existing data set and in both data sets, respectively, m*^(*NDC*)^ *and* 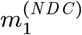 *is the number of test units and the number of alternative test units in the NDC*.

**Proof.** The statement of the theorem directly follows from Lemma 2 for test units in the existing data set. The number of test units in the NDC and not in the existing data set is 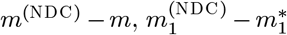 of which is alternative. Consequently, 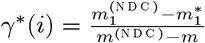 for test unit *i* that is not in the existing data set, because we have no prior information for this test unit from the existing data set.

##### Corollary 4

*Under the reasonable assumption that the concentration of the alternative test units is the same inside and outside the region covered by the existing data set, i.e*. 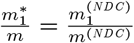, *we have that*

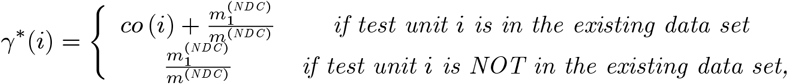

*and co* (*i*) *can be calculated as*

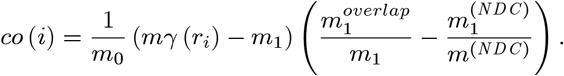

**Proof.** The statement of the corollary follows from Theorem 3 and from that 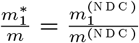 implies 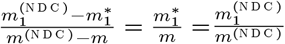.

##### Some properties of rank-based probability, γ, prior probability, γ*, and the contribution

First we derive formulas for rank-based probabilities that we will use to obtain formulas for prior probabilities and contribution.

###### Theorem 5

*Suppose that X*_1_,…,*X*_*m*_0__ *are identically (not necessarily independently) distributed random variables, representing the true null statistics, and suppose that Y*_1_,…, *Y*_*m*_1__ *are identically (not necessarily independently) distributed random variables, representing the true alternative statistics. Denote the kth largest random variable from* {*X*_1_,…, *X*_*m*_0__, *Y*_1_,…,*Y*_*m*_1__} *as Z_k_. Then for any fixed i, the probability that Y_i_ is the kth largest test statistic value, Z_k_, is*

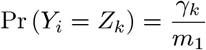

*and the probability that X_i_ is the kth largest test statistic value is*

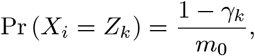

*where γ_k_ is the probability that Z_k_ is alternative*.

**Proof.**

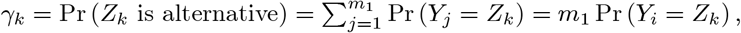

from which the first statement follows. The second statement can be proven similarly.

###### Corollary 6

*As a consequence of the above theorem we have that*

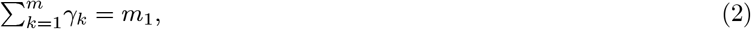

*where m = m*_1_ + *m*_0_.

**Proof.** For any fixed *i*, the events 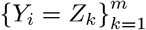 is a partition of the the probability space, we have that

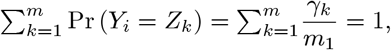

which implies the statement of the corollary.

###### Remark 7

*Note that the analogous statement in Corollary 6 for parametric calculation is not valid. That is if the parametric formula*

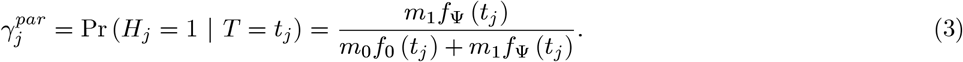

*is used to calculate the probability that a test unit is alternative in the existing data set, where f*_0_ *and f*_ψ_, *are the null and the alternative p.d.f., then* 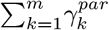 *may not be equal with m*_1_. *However, it is easy to see that*

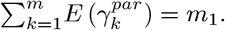

###### Lemma 8

*For any existing data set we have that*

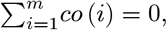

*where m is the number of test units in the existing data set.*

**Proof.**

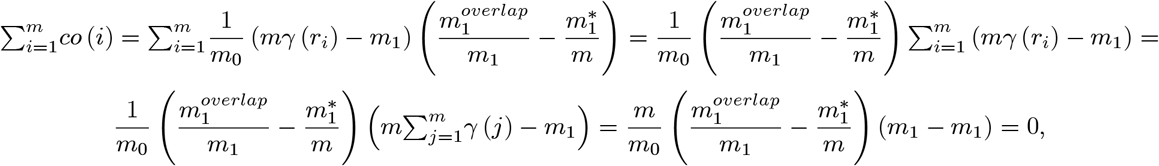

where we used (2).

###### Corollary 9

*If we have a single existing data set, then*

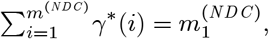

*where m*^(*NDC*)^ *and* 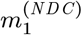 *is the number of test units and the number of alternative test units in the NDC, respectively. That is the sum of the prior probabilities is the same as the one without prior information.*

**Proof.** We have that

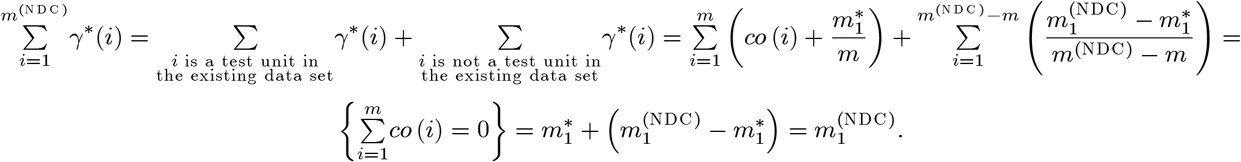

###### Claim 10

*We have that*

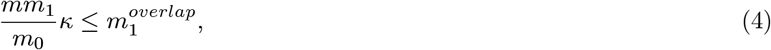

*and equality holds if and only if* 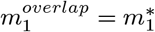.

**Proof.**

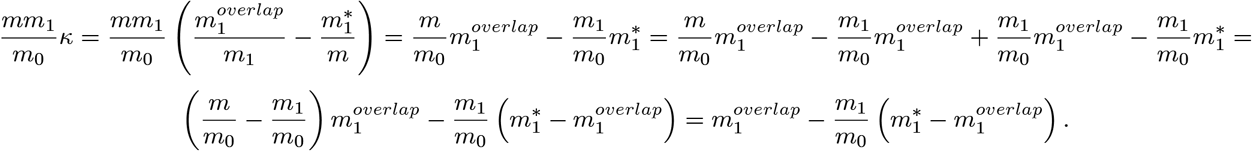

Clearly,

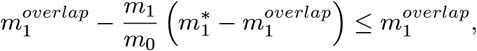

and equality holds if and only if 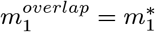.

##### Existing data sets with ties

Suppose we have *t* groups of test units and the ranks of all test units in a group are identical. Let *R_i_* be the number of test units whose rank is the *i*th largest one or smaller for *i* = 1,…, *t*, and let *R*_0_ = 0. Clearly, *R_i_ − R*_*i*−1_ is the number of test units in the ith group for all *i* = 1,…, *t*. Denote the probability that a test unit in the ith group is alternative in the existing data set as Γ_*i*_.

###### Claim 11

Γ*_i_ can be calculated by*

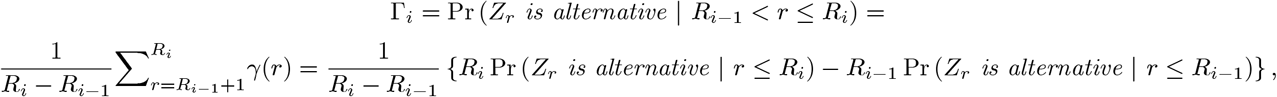

*where Z_r_ and γ*(*r*) *are defined above*.

###### Corollary 12

*Denote the probability that a test unit in the ith group is alternative in the NDC as* 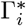. *Then we have that*

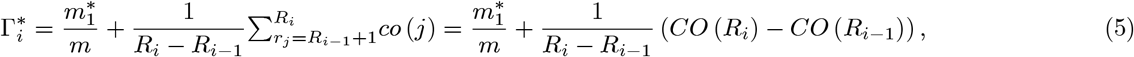

*where we define* 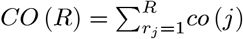.

**Proof.** We prove first that

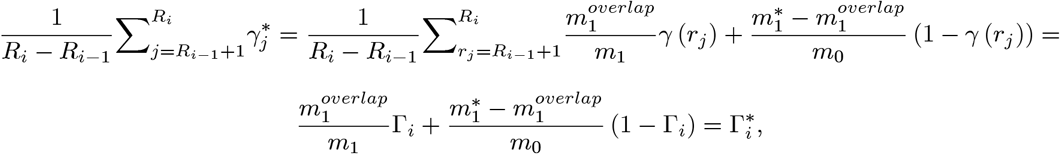

where the last step can be proved analogously to (1). Then it readily follows from Lemma 2 that

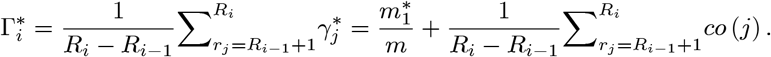

The statement analogous to the one in Corollary 6 is also valid:

#### Claim 13

*The sum of the prior probabilities will be m*_1_, *i.e*.

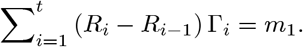

**Proof.**

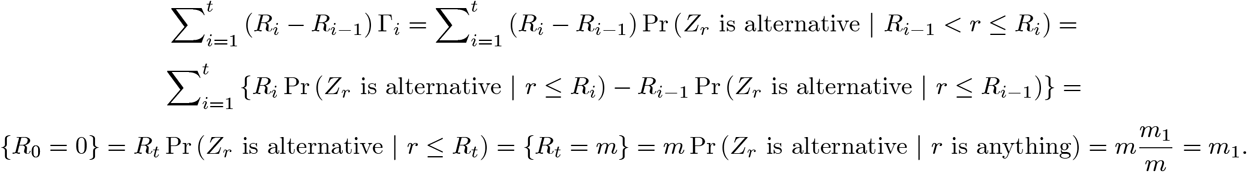

###### Claim 14

*Suppose that X*_1_,…, *X*_*m*_0__ *are i.i.d. random variables with F*_0_ *and Y*_1_,…, *Y*_*m*_1__ *are i.i.d. random variables with F*_ψ_. *Suppose that X*_1_,…, *X*_*m*_0__, *Y*_1_,…,*Y*_*m*_1__ *are independent, and denote the rth largest random variable among them as Z_r_. Then the probability that Y_s_ is in the ith (largest) group of test units is*

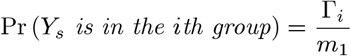

*and the probability that X_s_ is in the ith (largest) group of test units is*

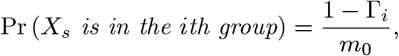

*where* Γ*_i_ is the probability that Z_k_ is alternative given it is in the ith group*.

**Proof.** The probability that *Y_s_* is in the ith (largest) group of test units is

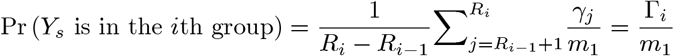

and the probability that *X_s_* is in the ith (largest) group of test units is

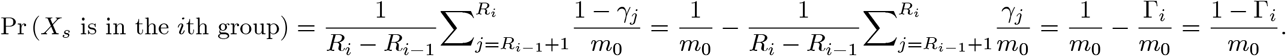

###### Remark 15

*Note that the distribution in Theorem (5) and that in Claim 14) is formally the same. As a result, for multiple instances we do not need to distinguish the case ties from that of no ties*.

#### Step 2: Combining the sets of prior probabilities into a single set of prior probabilities

##### Definition 16

*Suppose we have k existing data sets with (ranks of) test statistic values* 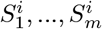. *The combined prior probability that a test unit is alternative in the novel data collection based on the information in the existing data sets is defined as*

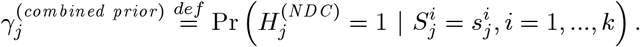

##### Theorem 17

*Suppose we have k existing data sets, and* 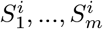 *are the test statistic values or the ranks of the test statistic values in the ith existing data set, i* = 1,…, *k, where some* 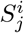 *may be missing. Denote the number of alternative test units in the ith existing data set as* 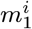. *Then we have that*

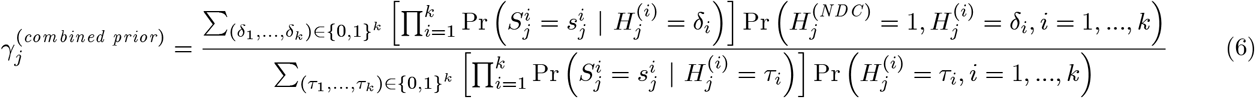

*where m*_1_ *and m*_0_ *is the number of alternative and null test units, respectively, in the novel data collection. Notation* {0, 1}*^k^ means all the 0-1 vectors of length k. Moreover*,

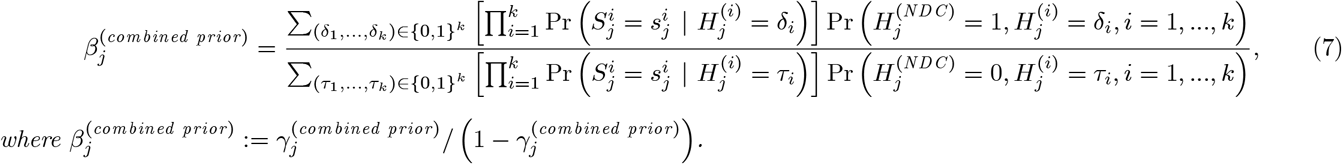

**Proof.** We have that

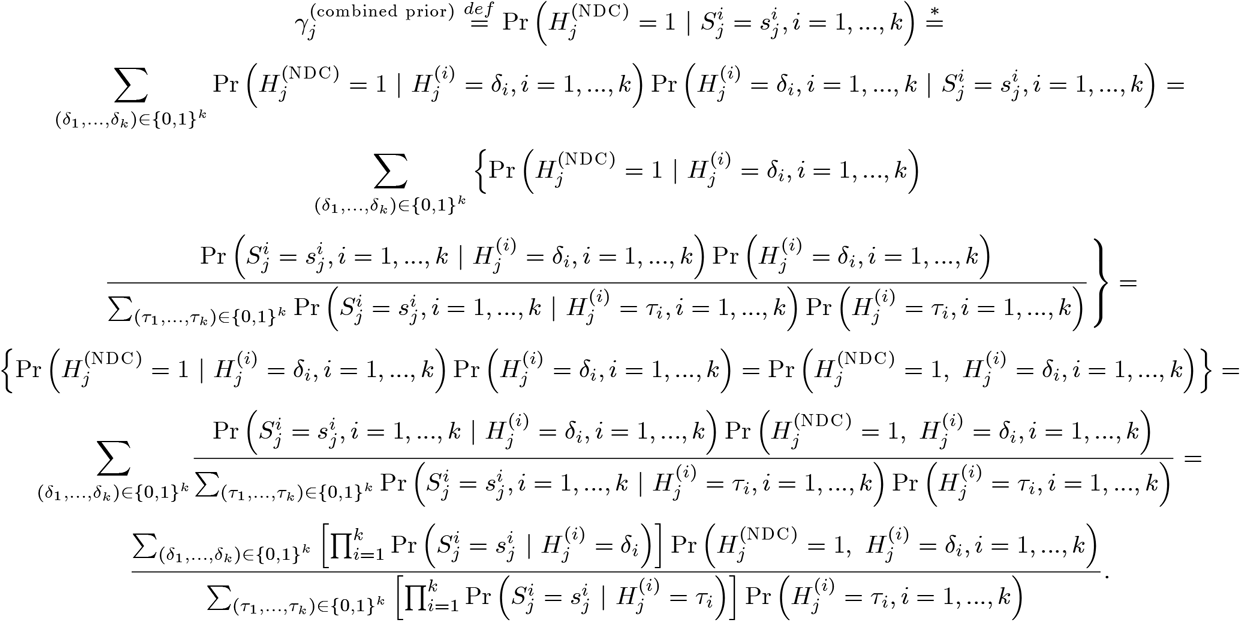

At ∗ we used that

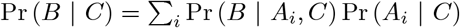

if *A_i_, i* = 1, 2… is a partition of the probability space. Moreover, we used the reasonable assumption that 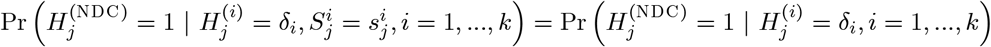.

As it is unknown in practice how the sets of alternative test units in the existing data sets and novel data collection overlap, the probabilities 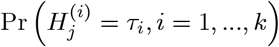 and 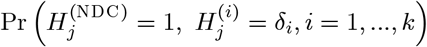 in (6) are unknown. Therefore, we need to use some mild assumptions to provide some practically useful and sufficiently accurate methods to combine prior probabilities from several existing data sets.

##### Theorem 18

*Suppose we have k existing data sets. Suppose that*

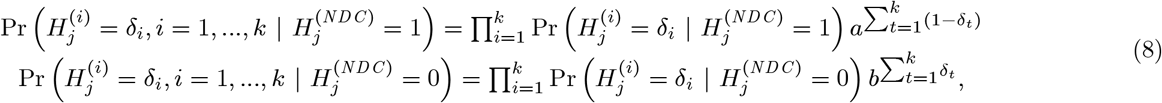

*hold for every* (*δ*_1_,…, *δ_k_*) ∈ {0, 1}^*k*^, *where* 0 ≤ *a, b* ≤ 1 *and we define* 0^0^ = 1. *Then*

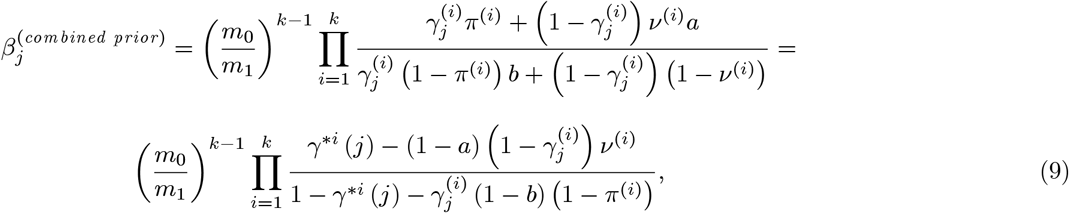

*where m*_1_ *is the number of alternative test units in the novel data collection, m*_0_ = *m − m*_1_,

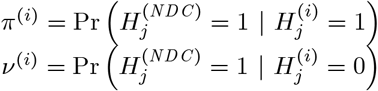

*for i* = 1,…, *k*,

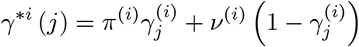

*is the prior probability that test unit j is alternative in the NDC based on the ith EDS (see (1)), and* 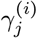 *is the probability that test unit j is alternative the ith EDS, i.e.*

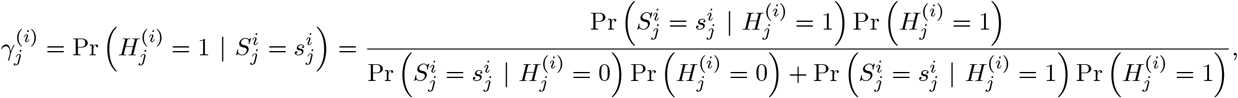

*where* 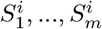 *are the test statistic values or the ranks of the test statistic values in the ith existing data set, i* = 1,…, *k*.

**Proof.** Applying criterion in (8) for the formula in (7) we obtain that

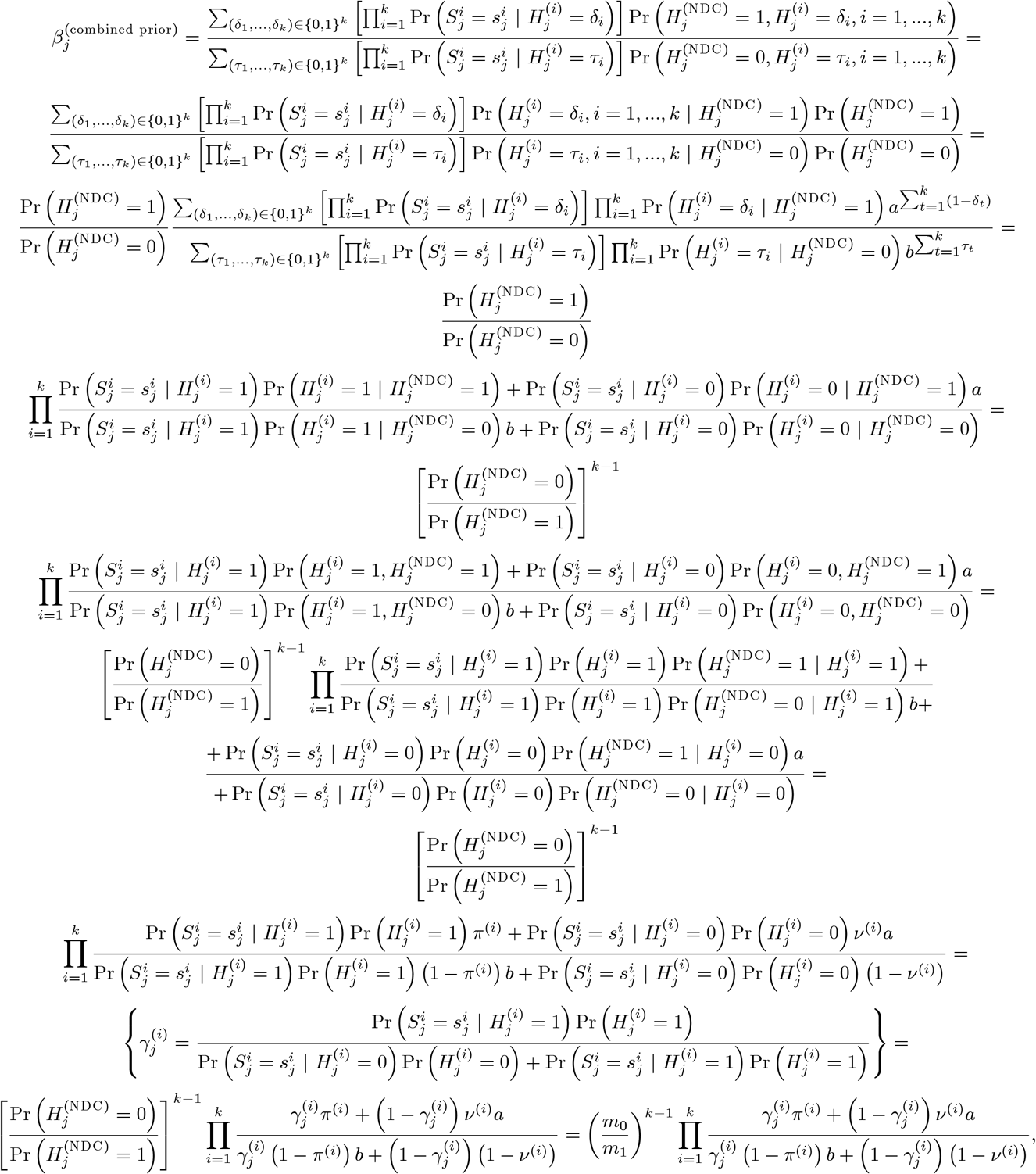

which proves the first equality in (9). Moreover, we have that

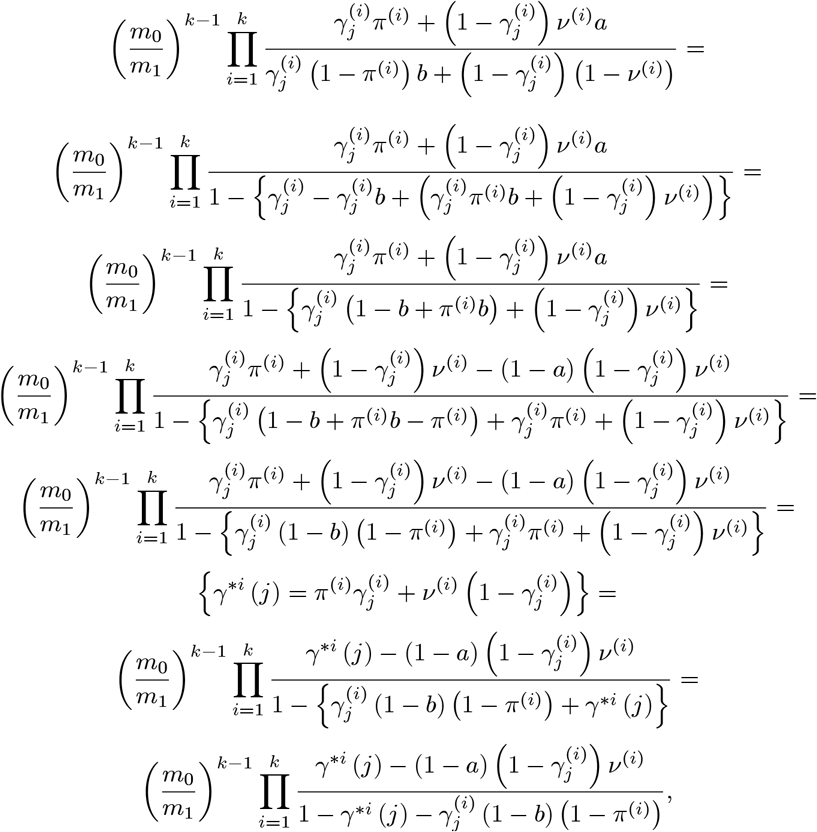

which proves the second equality in (9).

##### Corollary 19

*Suppose we have k existing data sets. Suppose that*

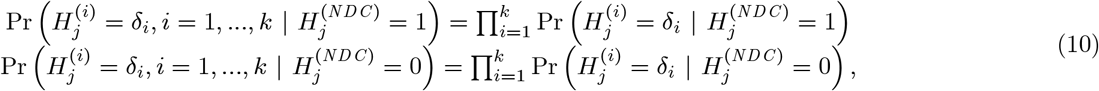

*for every* (*δ*_1_,…, *δ_k_*) ∈ {0,1}*^k^, then*

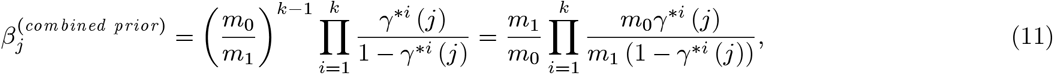

*where m*_1_ *is the number of alternative test units in the novel data collection, m*_0_ = *m − m*_1_, *and γ^*i^*(*j*) *is the prior probability that test unit j is alternative in the NDC based on the ith EDS*.

**Proof.** The condition in (10) is equivalent to (8) with *a = b* = 1. Therefore, substituting *a = b* = 1 in (9) we obtain that

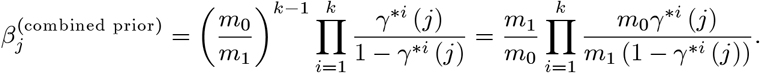

##### Corollary 20

*If*

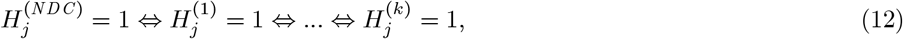

*then*

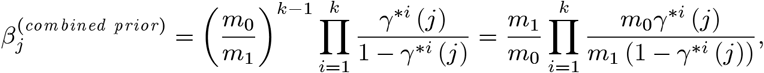

*where m*_1_ *is the number of alternative test units in the novel data collection, m*_0_ = *m − m*_1_, *and γ_*i_*(*j*) *is the prior probability that test unit j is alternative in the NDC based on the ith EDS.*

**Proof.** First we need to see that the condition in (12) is equivalent to (8) for *a = b* = 0. Indeed, substituting *a = b* = 0 in (8) we obtain that

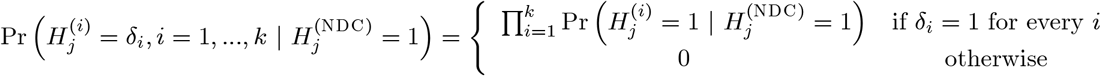

and

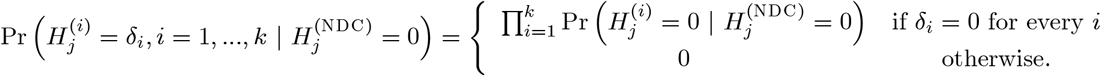

As

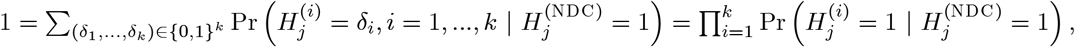

we have that 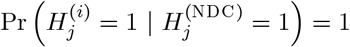 for every *i*, hence 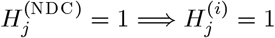 for every *i*. Similarly as

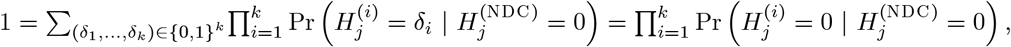

we have that 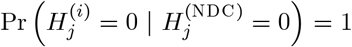 for every *i*, hence 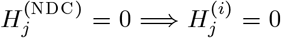 for every *i*. These two together implies 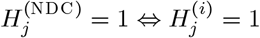 for every *i*.

Also the condition in (12) implies 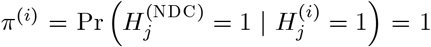 and 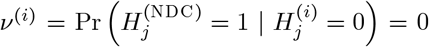. Therefore, substituting *a = b* = 0, *π*^(*i*)^ = 1 and *v*^(*i*)^ = 0 in (9) we obtain that

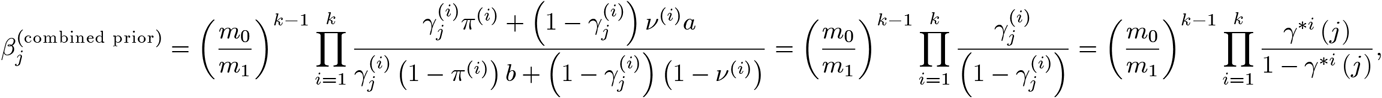

where the last equation holds because 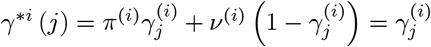, as *π*^(*i*)^ = 1 and *v*^(*i*)^ = 0. This completes the proof of the Corollary.

##### Remark 21

*Note that the conditions in (10) and (12) represent the two extrema of (8), and in both cases the combined odds can be calculated as*

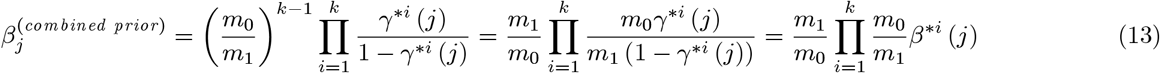

*where m*_1_ *is the number of alternative test units in the novel data collection, m*_0_ = *m − m*_1_, *γ^*i^*(*j*) *is the prior probability that test unit j is alternative in the NDC based on the ith EDS, and the odd β^*i^*(*j*) *is defined as β^*i^*(*j*) = *γ^*i^*(*j*) / (1 − *γ^*i^*(*j*)). *Moreover, the terms in the product in (9) can be approximated with β^*i^*(*j*) *even if (10) and (12) do not hold, suggesting that the formula in (13) is reasonable even for the general case. For the general formula (6) the structure of how the sets of test units alternative in the EDSs as well as the set of test units alternative in the NDC overlap each other need to be known, which may be difficult to estimate*.

##### Remark 22

*Recall that from (1) we have that*

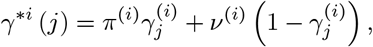

*where*

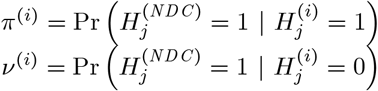

*for i* = 1,…, *k, (see (1)), and* 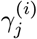 *is the probability that test unit j is alternative the ith EDS, i.e*.

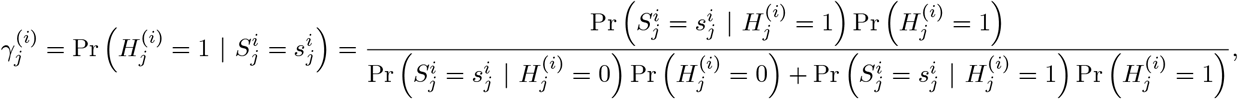

*where* 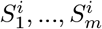 *are the test statistic values or the ranks of the test statistic values in the ith existing data set, i* = 1,…, *k*.

#### Step 3: Computing c*ℓ*TDR for each test unit

The compound *ℓTDR* (*cℓTDR*) of a test unit is defined as the posterior probability that the test unit is alternative in the novel data collection based on the information we have from the existing data sets and the novel data collection. The mathematical definition of the *cℓTDR* of a test unit is the following.

##### Definition 23

*The compound ℓTDR (cℓTDR) of test unit j is defined as*

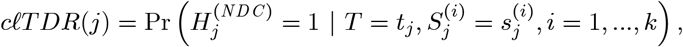

*where t_j_ is the observed test statistic value of test unit j in the novel data collection and* 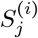 *is the test statistic value or the rank of the test statistic value of test unit j in the ith existing data set, i* = 1,…, *k*.

##### Claim 24

*The cℓ TDR of a test unit can be calculated as*

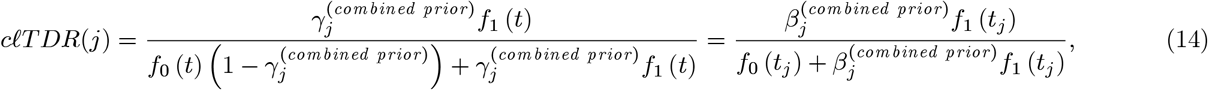

*where f*_0_ *and f*_1_ *is the null and alternative p.d.f. in the novel data collection, respectively*, 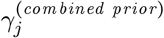 *is the combined prior probability (from the existing data sets) that test unit j is alternative in the novel data collection, and* 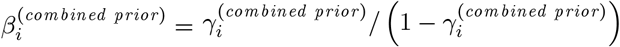.

**Proof.** We have that

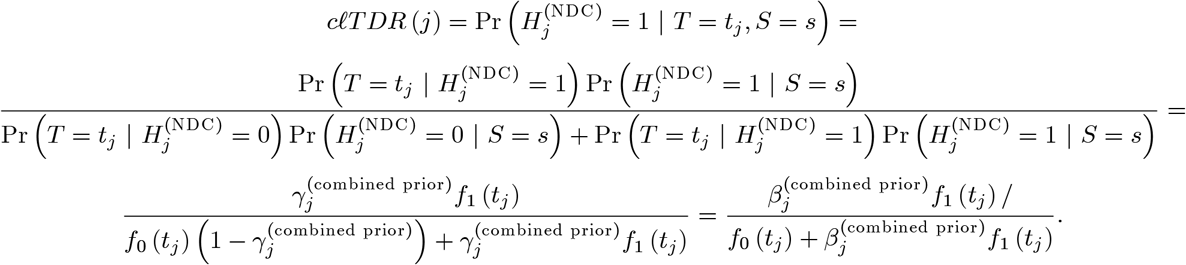

The estimator of the c*ℓ*TDR can be obtained by substituting 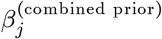, *f*_0_(*t*) and *f*_1_ (*t*) and with their estimates in (14).

### 1.2 Estimating the contributions

#### Theorem 25

*Denote the number of test units that are both in the NDC and the existing data set as m. For a positive integer M* ≤ *m and real d* ≥ 0 *we have that*

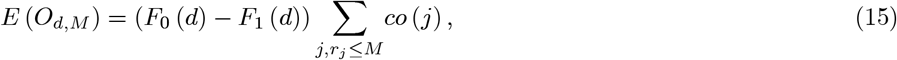

*where*

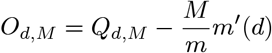

*and Q_d,M_ and m′*(*d*) *are defined as*

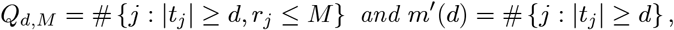

*and co*(*j*) *denotes the contribution of test unit j*.

**Proof.** Denote the set of test units alternative in the EDS and the ones alternative in the NDC as *E*_1_ and *N*_1_, respectively. Moreover, denote the set of test units that are null in the EDS and the ones null in the NDC as *E*_0_ and *N*_0_, respectively. We have that

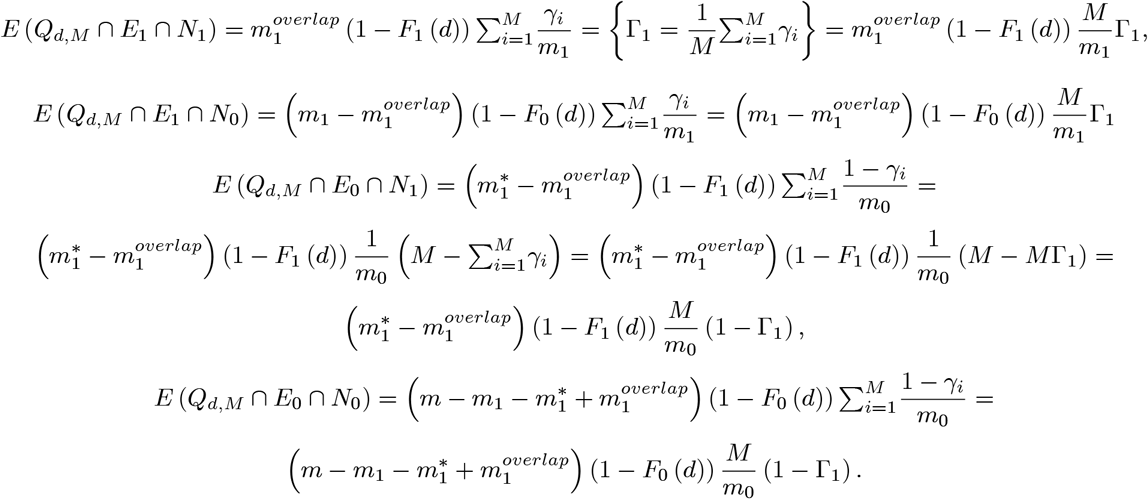

Therefore, we have that

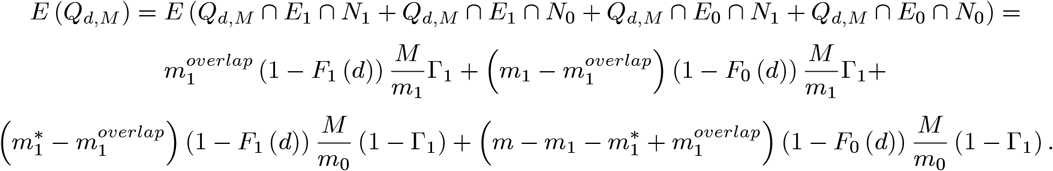

Moreover, we have that

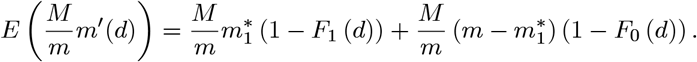

Combining the above two we obtain

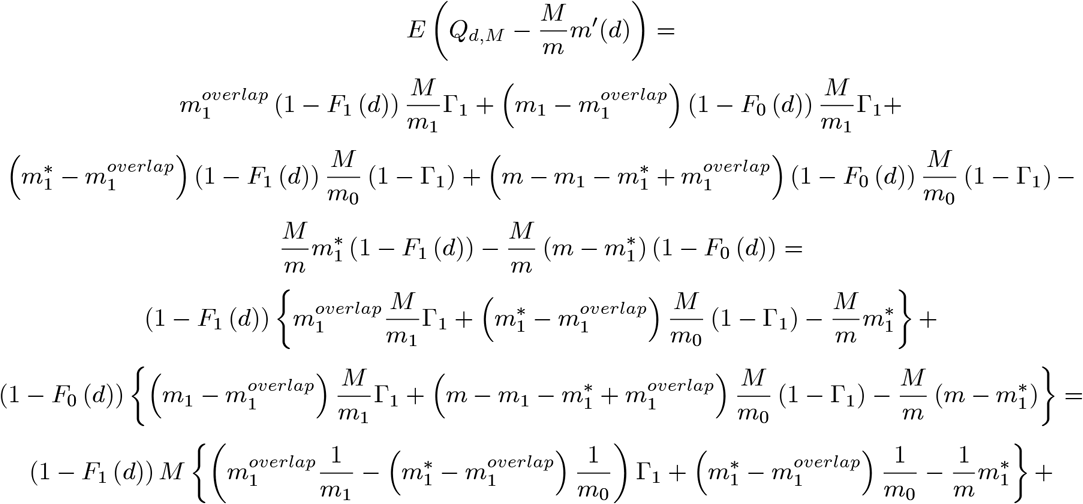

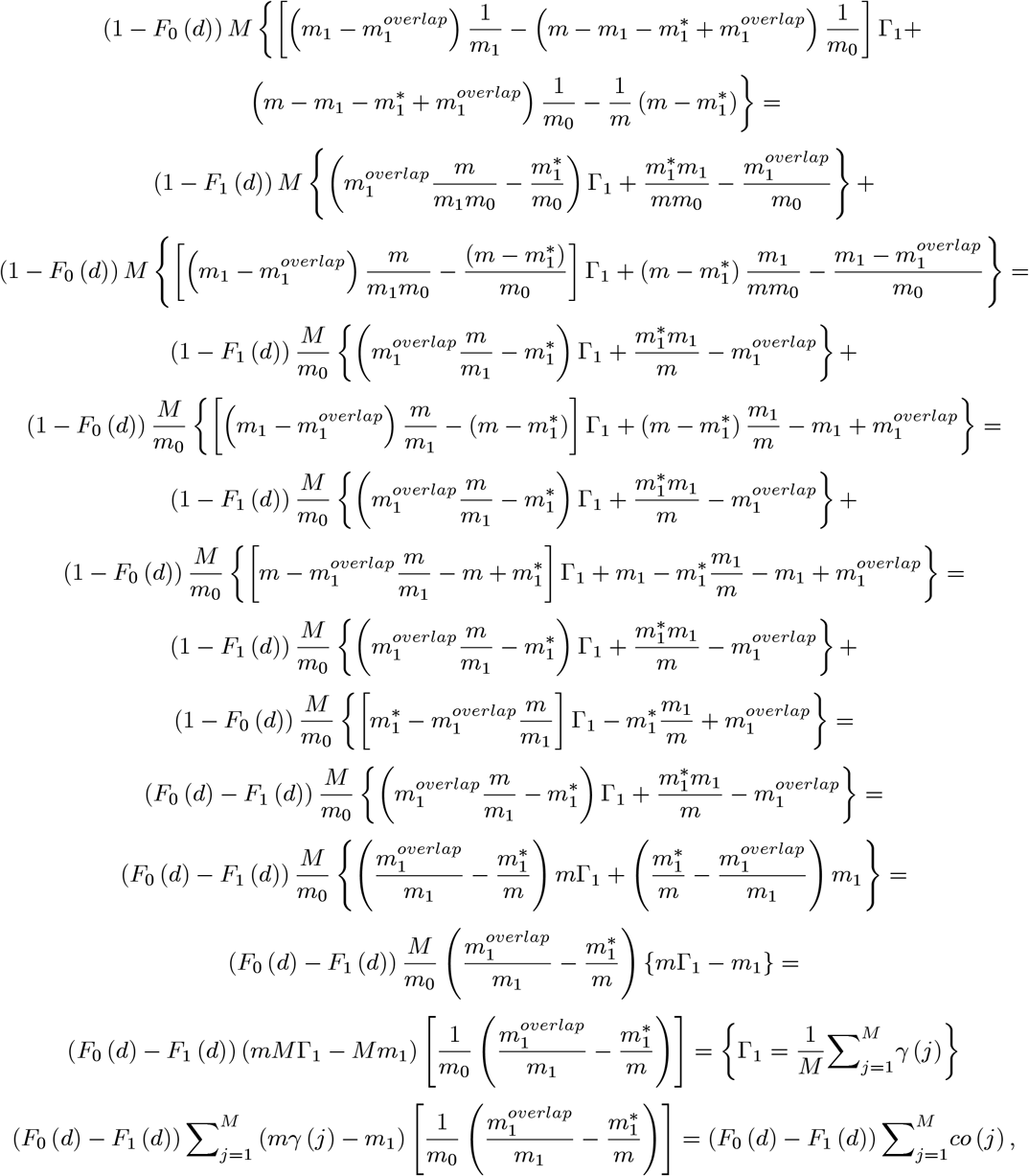

which concludes the proof of the theorem.

### 1.3 Smoother

In this section we present a method that smooths the 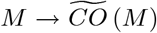 to a concave curve. In particular, we will choose the best fitting curve from the family of all *M → CO*(*M*) curves that can be attained by a suitable selection of parameters. We emphasize that our goal is not to find accurate estimates of the parameters, but to estimate the contributions accurately by a curve-fitting method. As a matter of fact, we may obtain similar curves by infinite many choices of parameter sets, thus, it is possible to approximate the true *M → CO*(*M*) curve very well while the parameters of the approximating curve may be far from the real parameter values.

From (15) and the definition of 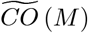 and the contribution we have that

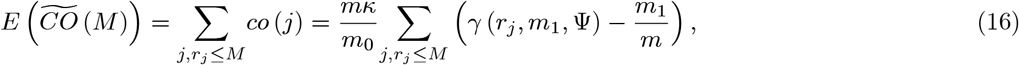

where the term on the right-hand side depends on three unknown parameters, *κ*, Ψ and *m*_1_, as *m*_0_ = *m − m*_1_. We will use the approximation

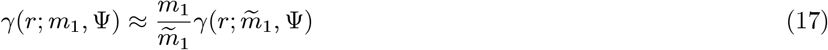

where 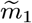 is an arbitrary choice, say our guess for the number of alternatives in the existing data set. By applying approximation (17) to the term on the right-hand side in (16), we obtain

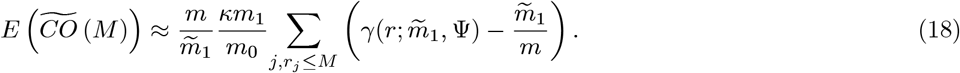

where the term on the right-hand side depends on only two unknown parameters, *κ*\* = *κm*_1_/*m*_0_ and Ψ, thus, it will be denoted as *CO*(*M; κ*\*, Ψ). By a curve-fitting method, we find 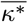 and 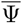 that provides the best fitting curve to 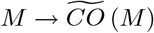 from the family of curves {*M → CO*(*M; κ*\*, Ψ), 0 < *κ*\*, Ψ}. Finally, 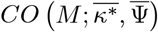 will be our estimate of the cumulative contribution, 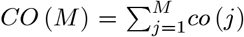. In the course of the algorithm, the term 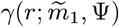 is computationally evaluated by the method described in section 1.3. In principle, we could use (20) or (21) to compute *γ*s, however, the numerical integration required is computationally very intensive, especially for large 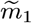, say 1,000 or larger. We remark that 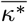 and 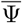 are not meant to be the estimator of *κ*\* and Ψ, as a matter of fact they may be far from the real *κ*\* and Ψ. Note that this is not a problem as long as 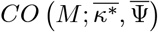 is an accurate estimator of the cumulative contribution. We remark that we have an estimator of Ψ, which plugged in (20) provides the estimator 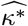 by a curve-fitting method. From (4) we have that 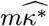 is a tight lower bound estimate of 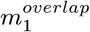.

#### Computation of the rank-based probability that a test unit has an effect in the existing data set, *γ*

In this subsection first we present an algorithm that computes the probability that a test unit is alternative in a data set, *γ*, utilizing the rank of the test statistic of the test unit in the data set. For completeness we will also present the formulas of *γ*, although using the algorithm is computationally much faster and not less accurate than applying numerical integrals to evaluate the formulas. We assume that the c.d.f. of the test statistic values is *F*_0_ and *F*_Ψ_ under the null and the alternative hypothesis, respectively. First we deal with the case when there are no ties in the ranking, then we deal with the case when there are ties in the ranking. Ties occur when the test units are sorted in a couple of categories, and our prior information does not distinguish between test units in the same category. For instance, we have only two categories if we have a candidate gene list, the test units that are on the list and those that are not on the list.

More formally, suppose that *X*_1_,…, *X*_*m*_0__ are identically distributed random variables with c.d.f. *F*_0_, and *Y*_1_,…, *Y*_*m*_1__ are identically distributed random variables with *F*_Ψ_. Suppose that *X*_1_,…, *X*_*m*_0__, *Y*_1_,…, *Y*_*m*1_ are independent, and denote the *r*th largest random variable among them as *Z_r_*. The probability that *Z_r_* is an alternative will be denoted as *γ*(*r*; *m*_1_, Ψ), or shortly *γ_r_*. In this section we give an algorithm and formulas in order to compute *γ_r_*.

#### Algorithmic computation of the probability that a test unit is alternative in the a data set based on its rank, *γ*

Based on the following theorem, 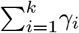 can be computed algorithmically, from which 7_i_s can readily be calculated.

##### Theorem 26

*For an integer k* = 1,…, *m we have that*

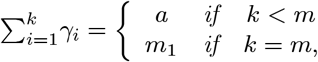

*where a is the solution of the equation*

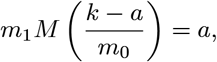

*where M*(*p*) *denotes the c.d.f. of the alternative p-values, i.e*. 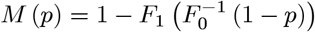. *Moreover, a* ∈ (max (*k − m*_0_, 0), min (*k, m*_1_)).

**Proof.** For *k = m*, the statement of the theorem follows from (2). Therefore, for the rest of the proof we can assume that *k < m.* By definition of *γ_i_*, we have that

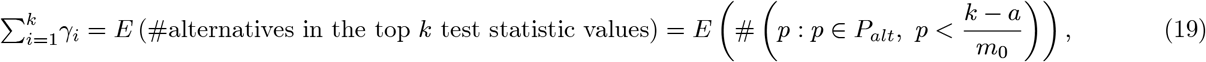

where *P_alt_* denotes the set of p-values of true alternative test units and *a* is selected in such a way that 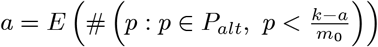. The equation in (2) holds because the expected number of alternative and null *p*-values smaller than 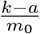 is *a* and *k − a*, respectively. Also, as we have that

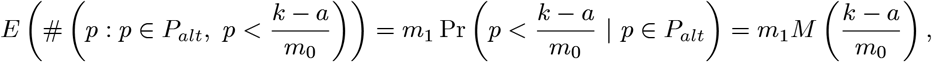

where *M*(*p*) denotes the c.d.f. of alternative *p*-values. In order to obtain 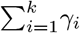 we need to solve

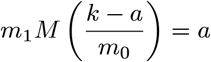

for *a*.

From (2) we have that

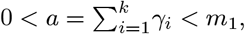

for *k* = 1,…, *m*−1, and, clearly 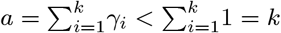, thus, *a* < min (*k, m*_1_). As 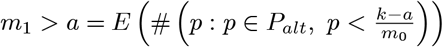, we have that 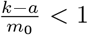, which implies *k − m*_0_ < *a*. As 0 < *a*, we have that max (*k − m*_0_, 0) < *a*, which completes the proof of the theorem.

#### Mathematical formulas of the probability that a test unit is alternative in the a data set based on its rank, *γ*

Now we give formulas of *γ* for the case when there are ties,and the case when there are no ties among the ranks.

##### Theorem 27

*Suppose that X*_1_,…, *X*_*m*_0__ *are i.i.d. random variables with F*_0_ *and Y*1_1_,…, *Y*_*m*_1__ *are i.i.d random variables with F*_Ψ_. *Suppose that X*_1_,…, *X*_*m*_0__, *Y*_1_,…, *Y*_*m*_1__ *are independent, and denote the rth largest random variable among them as Z_r_. Then the probability that Z_r_ is an alternative can be calculated as*

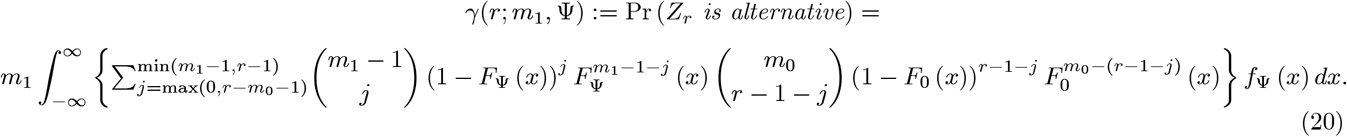

To prove the theorem we need the following lemma.

##### Lemma 28

*The marginal density function of Z_r_ intersected with the event that Z_r_ is alternative can be obtained as*

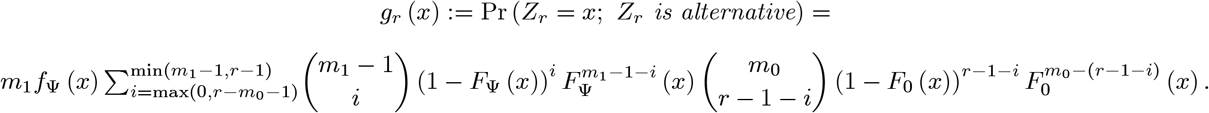

**Proof.**

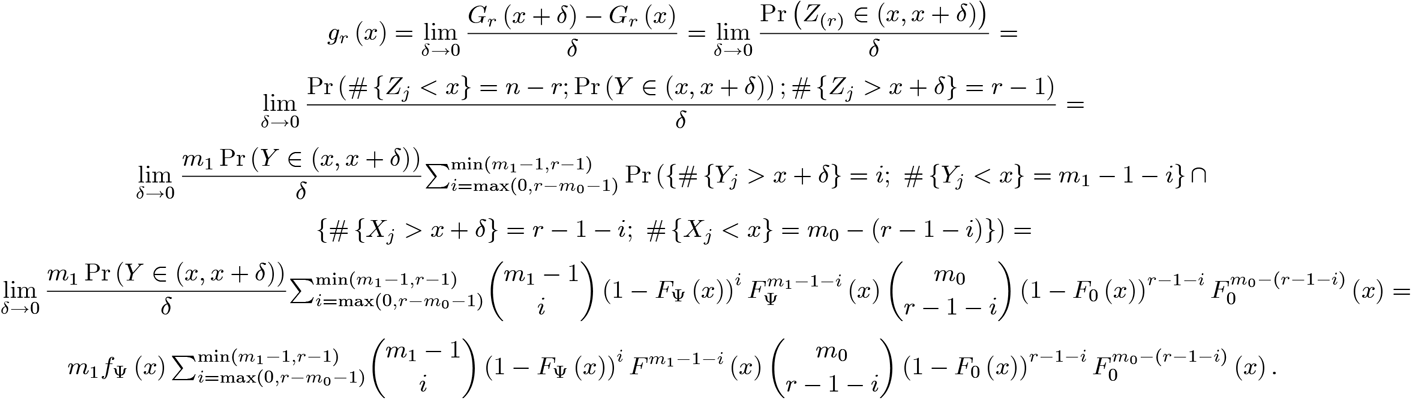

##### Proof of the Theorem.

Utilizing the lemma we have that

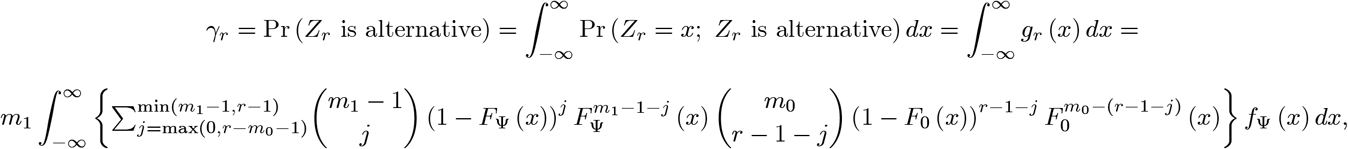

which completes the proof of the theorem.

##### Theorem 29

*With the reasonable assumption that m*_1_ ≤ *m*_0_ *we have that*

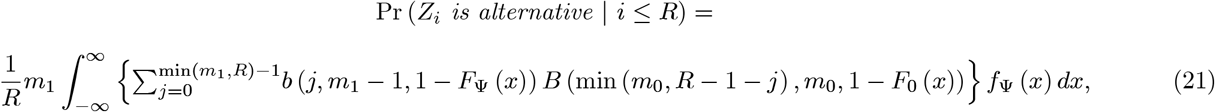

*where b* (*x,n,p*) *and B* (*x,n,p*) *are the binomial density and the distribution function values, respectively, at point x*.

**Proof.**

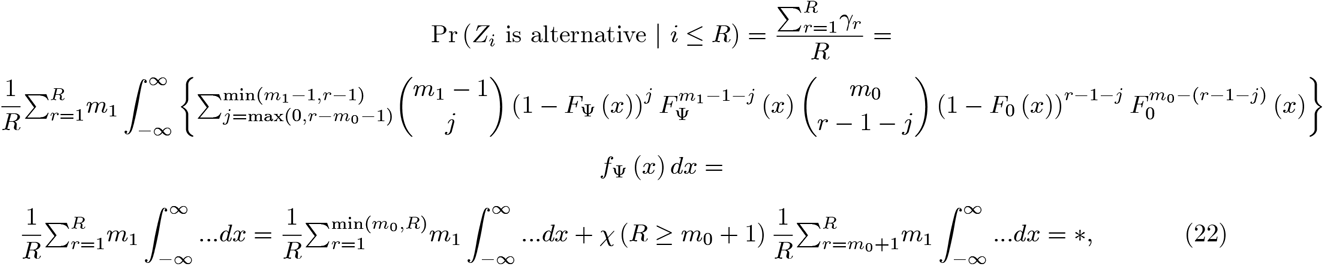

where *χ*(*A*) is the indicator function, i.e. *χ*(*A*) = 1 if *A* holds, and *χ*(*A*) = 0 otherwise. First we calculate the first sum. The rationale is that for *r* ≤ min (*m*_0_, *R*) we have max(0, *r − m*_0_ − 1) = 0.

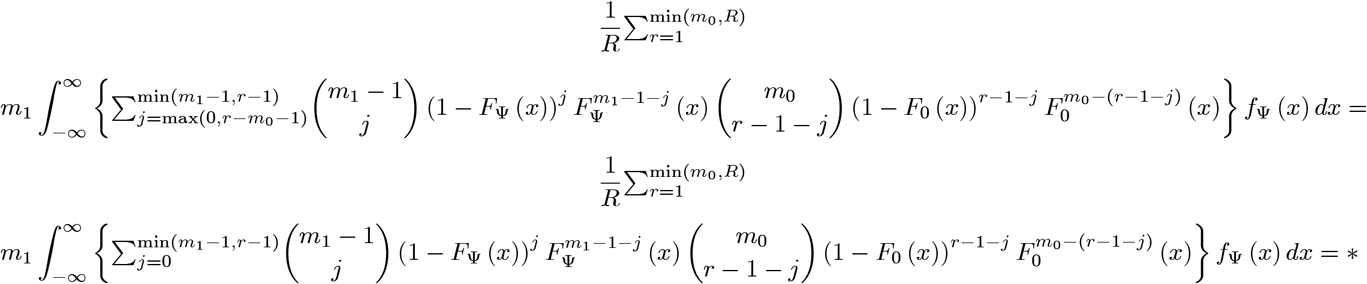

for the change of the sum

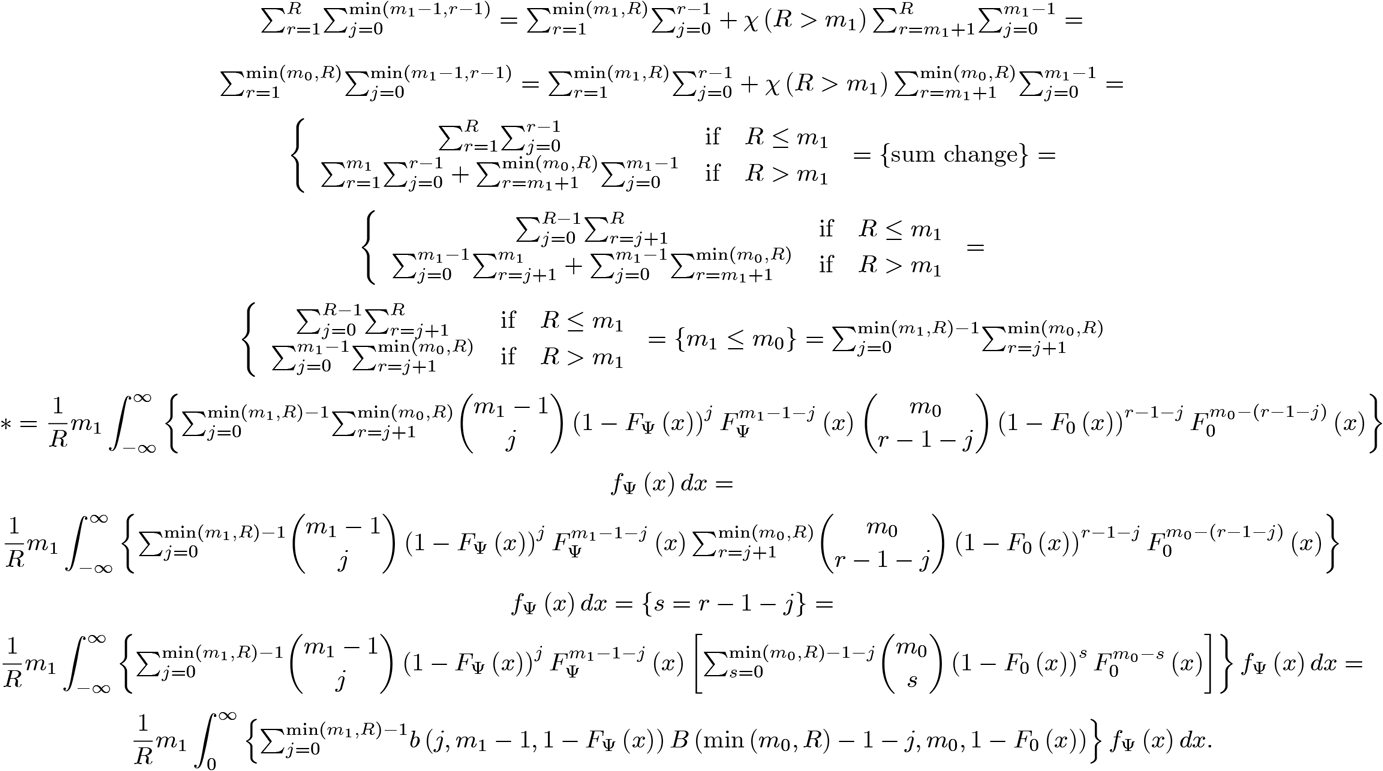

For the second sum in (22), as *r* ≥ *m*_0_ + 1 implies max(0; *r* − *m*_0_ − 1) = *r* − *m*_0_ − 1, we have that

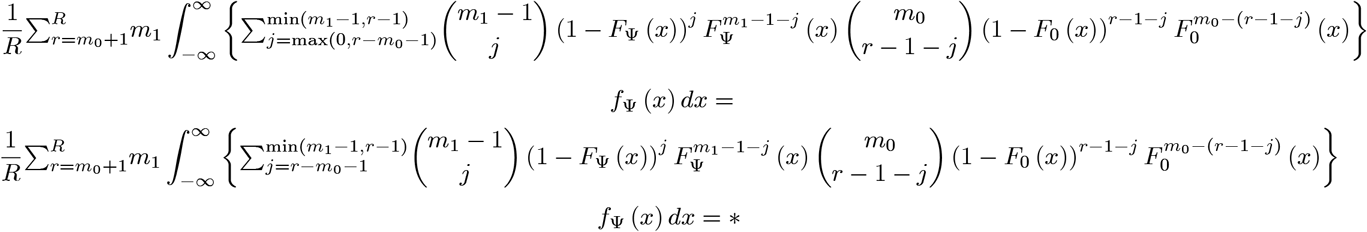

for the change of the sum we use that *m*_1_ ≤ *m*_0_

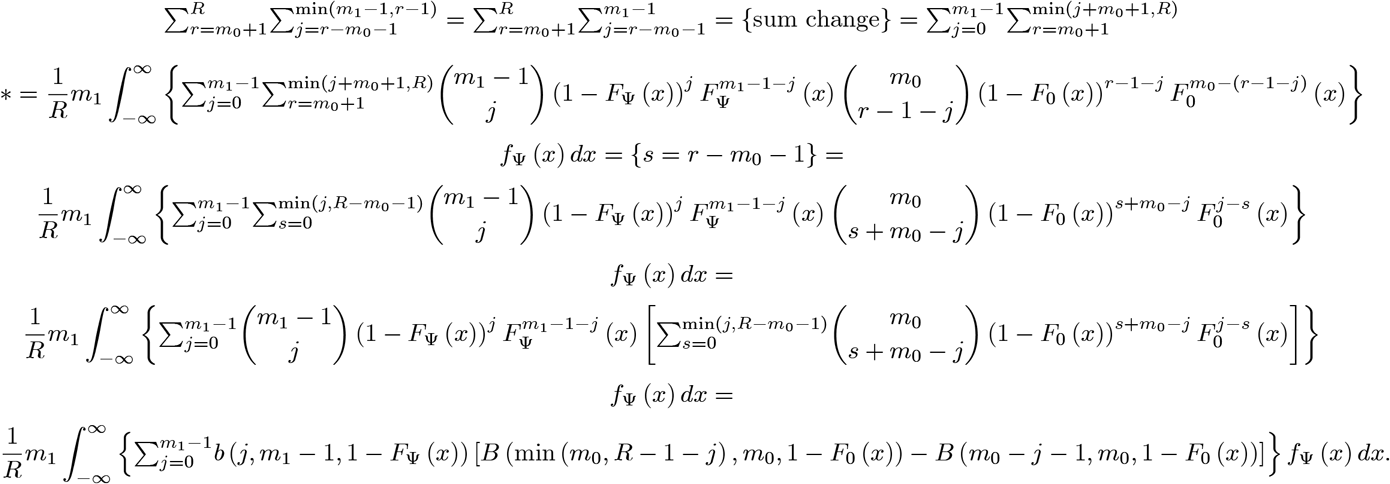

Putting the two sums together, from (22) we have that

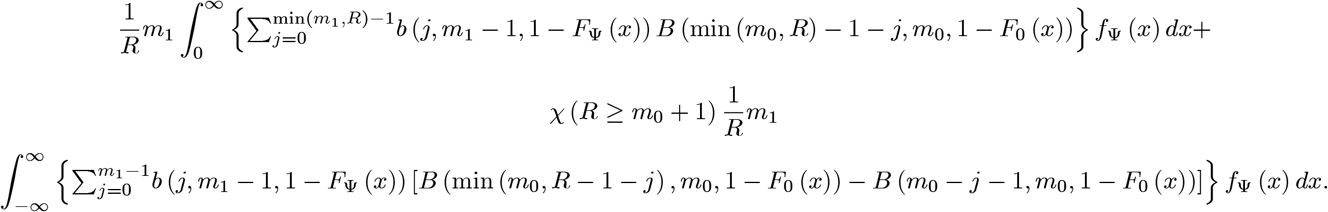

If *R* ≥ *m*_0_ + 1, then we have that

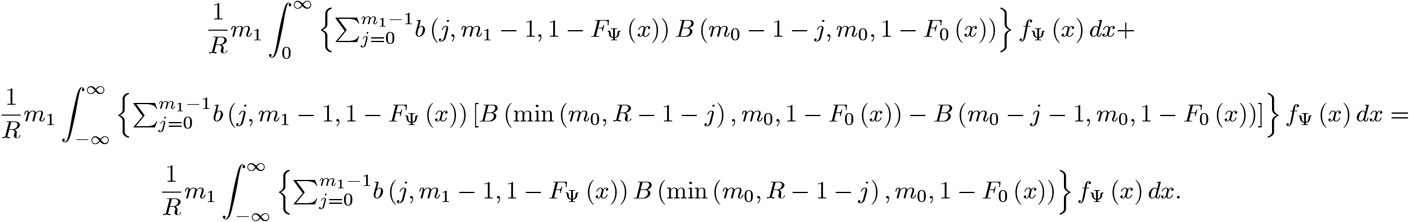

If *R* ≥ *m*_0_, then we have that

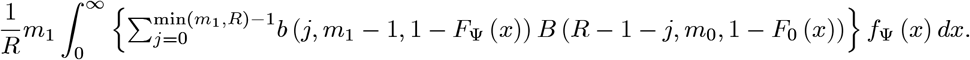

### 1.4 Overview of the algorithm that computes the estimates of *cℓTDR*

First we give the algorithm supposing that we have no ties in the ranks of the existing data sets, then we show how this needs to be modified for the existing data in which there are ties in the ranks.

1. First we need to estimate *F*_0_(*d*) and *F*_1_(*d*) in the novel data collection. (Note that here both *F*_0_(*d*) and *F*_1_(*d*) can be a mixture distribution, i.e. *F*_0_(*d*) and *F*_1_(*d*) merely denotes the null and alternative distribution in the novel data collection.)
2. We estimate the cumulative contribution, 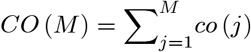 by first calculating

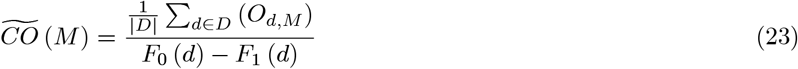

for *M* = 1,…, *m*, where *D* is a set of the positive real numbers, |*D*| is the number of elements in *D*. Then we apply a smoother method (see section 1.3) to fit 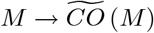 curve with a concave 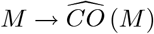 a concave function of *M*.
3. From 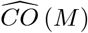 we calculate the estimator

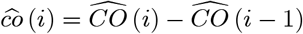

for *i* = 1,…, *m*, where we define 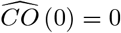.
4. Then 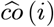 is used to calculate

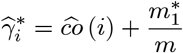

for every existing data set.
5. Then we use

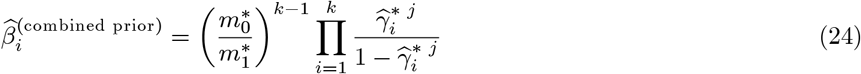

to calculate the estimates of combined prior odd for test unit *i* (see formula 13), where 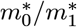 is the ratio of the null and alternative test units (markers) in the novel data collection, and 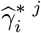 is the prior probability estimate of test unit *i* from the *j*th existing data set obtained in step 4.
6. The estimate of *cℓTDR* of test unit *i* will be calculated by

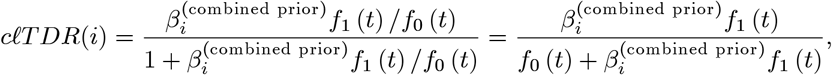

where *f*_0_ and *f*_1_ is the null and alternative p.d.f. in the novel data collection.

#### If there are ties among the ranks in an EDS, then Step 2 and 3 are modified for that EDS in the following way

Suppose we have *t* groups of test units and the ranks of all test units in a group are identical, but they are different across the groups. Let *R_j_* be the number of test units whose rank is the jth smallest one or smaller than that for *j* = 1,…, *t*. Then we calculate 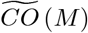 only for *M = R*_1_,…, *R_t_* in Step 2, and in Step 3 the estimator 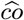 is obtained as

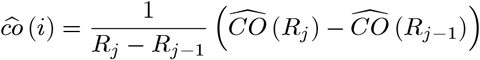

for every test unit *i* in group *j, j* = 1,…, *t*, where we define *R*_0_ = 0 and 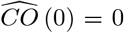 for the sake of simplicity (see (5) for justification).

We remark that in order to decrease computational burden, this modification in Step 2 and 3 can be used for the case of no ties as well. Note that in case of no ties, the choice of *R*_1_ <… < *R_t_* is not determined by the groups of ties in the rank of test units in the EDS, but the desired accuracy of the contribution estimator.

## Notes

### Competing Interest Statement

The authors have declared no competing interest.

